# Prenatal inflammation reprograms hyperactive ILC2s that promote allergic lung inflammation and airway dysfunction

**DOI:** 10.1101/2023.11.20.567899

**Authors:** Diego A. López, Aleah Griffin, Lorena Moreno Aguilar, Cassandra-Deering Rice, Elizabeth J. Myers, Kristi J. Warren, Robert Welner, Anna E. Beaudin

## Abstract

Allergic asthma is a chronic respiratory disease that initiates in early life, but causal mechanisms are poorly understood. Here we examined how prenatal inflammation shapes allergic asthma susceptibility by reprogramming lung immunity from early development. Induction of Type I interferon-mediated inflammation during development provoked expansion and hyperactivation of group 2 innate lymphoid cells (ILC2s) seeding the developing lung. Hyperactivated ILC2s produced increased IL-5 and IL-13, and were associated with acute Th2 bias, eosinophilia, and decreased Tregs in the lung. The hyperactive ILC2 phenotype was recapitulated by adoptive transfer of a fetal liver precursor following exposure to prenatal inflammation, indicative of developmental programming. Programming of ILC2 function and subsequent lung immune remodeling by prenatal inflammation led to airway dysfunction at baseline and in response to papain, indicating increased asthma susceptibility. Our data provide a link by which developmental programming of progenitors by early-life inflammation drives lung immune remodeling and asthma susceptibility through hyperactivation of lung-resident ILC2s.

**One Sentence Summary:** Prenatal inflammation programs asthma susceptibility by inducing the production of hyperactivated ILC2s in the developing lung.

## INTRODUCTION

Allergic asthma (AA) is a chronic respiratory illness that causes significant morbidity in an estimated 25 million Americans. AA often initiates in early life, and childhood asthma is the leading cause of chronic disease in children and causes most childhood hospitalizations (*1, 2*). Respiratory infections in early life are strongly associated with an increased risk for childhood asthma (*3–7*). The most common respiratory early-life viruses are human rhinoviruses (HRV) and respiratory syncytial virus (RSV), both single-stranded RNA viruses that invoke an initial systemic Type I interferon response (*8, 9*). Although early-life infection is among the strongest predictors of AA susceptibility, causal mechanisms are debatable – it is unclear whether early-life infection exposes genetic susceptibility or whether infection fundamentally impairs lung immunity; this distinction has been difficult to dissociate in human studies. The developing immune system is more plastic and sensitive to inflammation (*10, 11*), and there is an increasing need to understand how antenatal infection and associated inflammation affect the developing immune system and how this confers disease susceptibility.

AA is associated with T-helper type 2 (Th2) cell biased responses further triggered by exposures to common environmental allergens including house dust mite (HDM), animal dander, and pollen. Whereas the adult Th2 response in asthma has been well-elucidated (*12*), neonatal lung immunity is considered “immature” due to synergy between the lack of antigen exposure and a limited adaptive immune response (*13, 14*). Recent work has identified group 2 innate lymphoid cells (ILC2s), a class of innate-like lymphocytes, as critical mediators of asthma pathogenesis in mouse models (*15–19*). ILC2s arise in the developing lung during fetal development (*20*) and achieve maximal expansion prior to maturation of the adaptive immune response (*21, 22*). Unlike CD4+ T-helper cells, the adaptive immune counterpart of ILC2s, the latter lack antigen-specific receptors. Instead, ILC2s are activated by alarmin signals in their tissue microenvironment, such as epithelial-derived IL-25 and IL-33, resulting in potent IL-5 and IL-13 cytokine production (*23*). In mouse models of AA, ILC2-derived IL-5 and IL-13 have been shown to promote airway hyperactivity via eosinophil infiltration, mucus production, and epithelial cell proliferation (*15, 19, 24–26*). ILC2s can promote AA development in the absence of mature B- and T-cells in Rag mice (*17*), and depletion of ILC2s can ameliorate airway inflammation in response to allergen challenge (*16, 19, 27, 28*). Importantly, ILC2s have also been identified in humans and are increased in the airways of asthmatic patients, including children (*29, 30*). However, the role of ILC2s as early drivers of AA susceptibility in response to prenatal inflammation remains untested.

In the bone marrow (BM), ILC2 development is initiated through the orchestrated expression of transcription factors TCF-1, GATA3, Id2, and PLZF that facilitate commitment to the ILC lineage from the common lymphoid progenitor (CLP) (*31–34*). Putative ILC precursors have also been identified in the fetal liver (*35, 36*), and mature ILC2s are present in the fetal lung during late embryogenesis where they expand postnatally and establish tissue-residency (*20, 37*). Recent fate-mapping experiments have demonstrated “layered” immune development of ILC2s in the lung: the initial wave of fetal ILC2s is replaced by neonatal ILC2s that are eventually replaced by a later wave of ILC2s in the adult (*20*), although the sources for different waves have not been investigated. Tissue-resident cells that seed developing tissues are increasingly implicated in regulating tissue immunity and homeostasis. A prime example includes tissue-resident macrophages in the lung that phagocytose pathogens and clear mucus for gas exchange (*38*). However, whether tissue-resident cells like lung ILC2s play a direct role in educating tissue immunity during early life has not been directly investigated.

Here we investigated the role of prenatal inflammation in driving asthma susceptibility by programming tissue-resident ILC2s that shape lung immunity during development. We recently demonstrated that prenatal inflammation shapes the production and function of innate-like lymphocytes postnatally by specifically activating a lymphoid-biased fetal hematopoietic progenitor (*39*). We proposed that reprogramming the function and output of tissue-resident immune cells that seed developing tissues during early life by prenatal inflammation would have fundamental consequences for tissue immunity, tissue function, and disease susceptibility.

## RESULTS

### Prenatal inflammation generates expanded and hyperactivated lung ILC2s in offspring

To investigate how prenatal inflammation impacts group-2 innate lymphoid cells (ILC2s) in the lung, we profiled changes to ILC2 establishment in offspring from postnatal day (P)3 through adulthood using a model of prenatal inflammation induced by administration of 20mg/kg poly(I:C) at embryonic day (E)14.5 (**Fig. 1)**. ILC2s (CD45+lineage-KLRG1+Sca1+Thy1+ST2+; **Fig. 1A**), were significantly expanded in response to prenatal inflammation as early as P9 and remained robustly expanded at P14 and into adulthood (10 weeks) (**Fig. 1B**). To assess the mechanism driving ILC2 expansion, we measured proliferation of ILC2s from P4 through P14. Lung ILC2s exhibit maximal proliferation during the first two weeks of life and peak in cellularity at P14 (*20, 21*). In response to prenatal inflammation, ILC2s exhibited heightened proliferative capacity at P4 and P9, based on Ki67 expression, that then normalized by P14 (**Fig. 1C-E**).

**Fig. 1:**
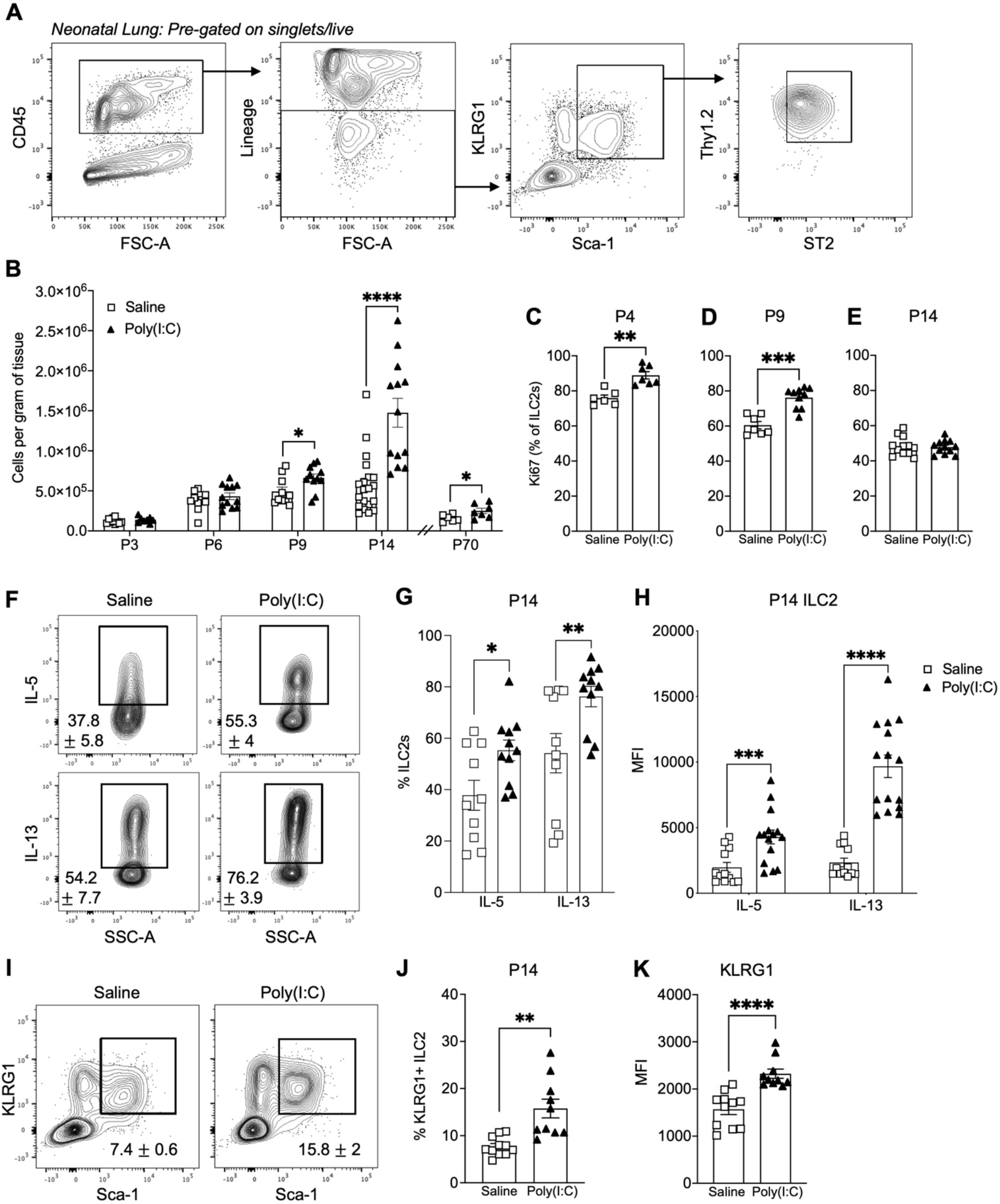
Prenatal inflammation induces expansion and hyperactivation of neonatal lung ILC2s. **A)** Representative flow cytometric gating of lung ILC2s from postnatal day (P)3-P70, pre-gated on Singlet and Live cells. **B)** Quantification of lung ILC2s (**Fig. 1A**) in offspring from P3-P70 after saline or poly(I:C) (20mg/kg) at embryonic day (E)14.5 **C-E)** Frequency of Ki67+ lung ILC2s at P4 **(C)** P9 **(D)** and P14 **(E)** in saline or poly(I:C) treated offspring **F-G)** Representative FACS plots **(F)** and frequency **(G)** of IL-5 and IL-13 positive ILC2s after *ex-vivo* stimulation with PMA + Ionomycin in P14 offspring after saline or poly(I:C) at E14.5. **H)** Mean Fluorescence Intensity (MFI) of IL-5 and IL-13 expression after *ex-vivo* stimulation with PMA + Ionomycin in lung ILC2s at P14 after saline or poly(I:C) at E14.5. **I-J)** Representative FACS plots **(I)** and frequency **(J)** of KLRG1+Sca1+ ILC2s at P14 after saline or poly(I:C) at E14.5. **K)** MFI of KLRG1 expression in ILC2s at P14 after saline or poly(I:C) treatment at E14.5. Data represent average ± SEM representing at least two independent experiments per condition. *p ≤ 0.05, **p ≤ 0.01, ***p ≤ 0.001, ****p ≤ 0.0001.

ILC2s are potent producers of IL-5 and IL-13 cytokines upon immune activation (*15, 19*). We next determined whether ILC2 expansion in the perinatal period was accompanied by alterations to their function. As we observed the most robust expansion in ILC2s at P14 (**Fig. 1B**), we assessed differential IL-5 and IL-13 production in P14 ILC2s isolated from offspring exposed to saline or prenatal inflammation after stimulation with PMA/ionomycin *ex vivo*. Prenatal inflammation robustly increased IL-5 and IL-13 production from ILC2s compared to saline controls (**Fig. 1F-H**). Both the frequency of IL-5- and IL-13-positive ILC2s (**Fig. 1G)** as well as IL-5 and IL-13 production in ILC2s (**Fig. 1H**) increased in response to prenatal inflammation. Concomitant with increased type-2 cytokine production, we also observed a higher frequency of ILC2s expressing killer cell lectin-like receptor G1 (KLRG1) (**Fig. 1J**) and greater expression of KLRG1 (**Fig. 1I, K**), which correlates with ILC2 activation (*40*). Thus, prenatal inflammation regulates neonatal lung ILC2 development and immune function by increasing their proliferative capacity and generating inflammatory and hyperactive ILC2s.

### Prenatal inflammation programs ILC2 hyperactivation from fetal precursors

We next investigated the origin of expanded and hyperactive ILC2s during the perinatal period. We recently demonstrated that poly(I:C)-induced prenatal inflammation specifically activates fetal lymphoid-biased developmentally-restricted hematopoietic stem cells (drHSCs) (*41*), resulting in expanded and hyperactivated innate-like B-1 B cells postnatally (*39*). To address if prenatal inflammation induced by poly(I:C) programs differential ILC2 hyperactivation at the progenitor level, we performed adoptive transfer experiments of drHSCs following induction of prenatal inflammation at E14.5. We crossed FlkSwitch male mice (*41–43*) to WT females and transplanted 200 fetal liver GFP+ drHSCs one day after saline or poly(I:C) treatment and analyzed donor-derived ILC2 progeny 4 weeks post-transplantation (**Fig. 2A**). Upon *ex-vivo* stimulation, a greater frequency of GFP+ ILC2s expressed both IL-5 and IL-13 cytokines in recipients of poly(I:C)-treated compared to saline-treated drHSCs (**Fig. 2B**). Furthermore, expression of IL-5 was also higher in recipients of poly(I:C)-exposed GFP+ drHSCs as compared to saline, as determined by MFI (**Fig. 2C-D**). These data demonstrate that prenatal inflammation programs increased production of IL-5 and IL-13 in mature lung ILC2s at the fetal HSC stage in a cell-intrinsic manner.

**Fig. 2:**
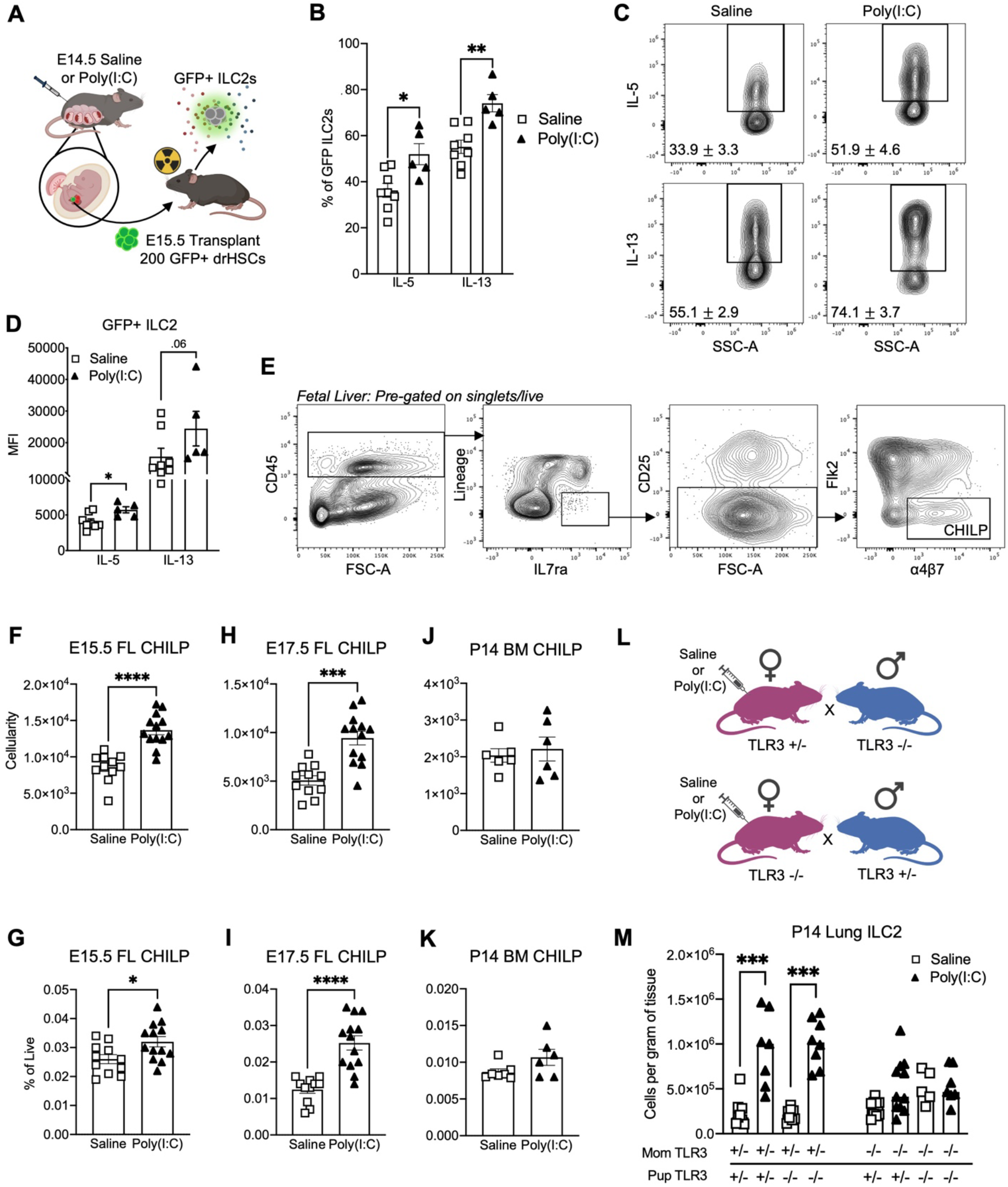
Prenatal inflammation programs ILC2 hyperactivation and expands ILC progenitors during fetal development. **A)** Schematic of sort and transplant of E15.5 fetal liver (FL) GFP+ developmentally-restricted HSCs (drHSC) following saline or poly(I:C) at E14.5. **B)** Frequency of GFP+ ILC2s expressing IL-5 or IL-13 after *ex-vivo* stimulation with PMA+ Ionomycin from saline or poly(I:C) treated GFP+ drHSC transplant recipients **C-D**) Representative FACS plots **(C)** and MFI **(D)** of IL-5+ or IL-13+ ILC2s from saline or poly(I:C) treated GFP+ drHSC transplant recipients. **E)** Representative FACS plots depicting gating strategy for CHILPs in the fetal liver (FL) at E17.5 **F-I)** Quantification **(F, H)** and frequency **(G, I)** of FL CHILP cells (Fig. 2E) at E15.5 (**F,G**) or E17.5 (**H, I**) following saline or poly(I:C) at E14.5 **J-K)** Quantification **(J)** and frequency **(K)** of bone marrow (BM) CHILPs in P14 offspring following saline or poly(I:C) at E14.5. **L)** Breeding schematic used to generate intercrossed TLR3 KO mice. Pregnant females were treated with saline or poly(I:C) (20mg/kg) at E14.5. **M)** Quantification of ILC2s in P14 offspring from intercrossed TLR3 KO mice after saline or poly(I:C) at E14.5. For all experiments, data represent average ± SEM representing at least two independent experiments per condition. *p ≤ 0.05, **p ≤ 0.01, ***p ≤ 0.001, ****p ≤ 0.0001.

Putative ILC progenitors have been identified at fetal and adult timepoints (*31, 35, 36, 44*). We profiled changes to an immediate upstream precursor of the ILC2 lineage, the common helper-like innate lymphoid progenitor (CHILP, Lin-IL7rα+Flk2-α_4_β_7_+CD25-) in the fetal liver one- and three-days post poly(I:C) injection (**Fig 2E**). Fetal liver CHILPs were robustly expanded by cellularity and frequency at embryonic day (E)15.5 and E17.5 (**Fig. 2F-I**). However, by P14, these changes within the fetal liver did not translate into persistent expansion of CHILPs by cellularity or frequency in BM of offspring (**Fig. 2J-K**). Importantly, mature ILC2s are not detectable in the fetal lung until E17.5 (*20*), suggesting that prenatal inflammation acts by directly expanding ILC progenitors rather than on the mature ILC2s themselves.

To further investigate how prenatal inflammation induced by poly(I:C) drove ILC2 expansion, we inter-crossed mice heterozygous or homozygous for deletion of TLR3, the receptor for poly(I:C), to disentangle the maternal and fetal response to poly(I:C) (**Fig. 2L**). When TLR3 signaling was deleted on the fetal side, but maternal TLR3 signaling was intact, we observed a significant expansion of ILC2s in the neonatal lung (**Fig. 2M**). This expansion was maintained regardless of partial or complete deletion of TLR3 signaling on the fetal side. In contrast, deletion of TLR3 on the maternal side completely ablated the effect of prenatal inflammation on the postnatal expansion of ILC2s in offspring (**Fig. 2M**). These data demonstrate that postnatal expansion of lung ILC2s requires poly(I:C) induced prenatal inflammation on the maternal side and does not depend directly on fetal TLR3 stimulation. Together, these data demonstrate that prenatal inflammation instructs fetal ILC precursors to expand, proliferate and generate hyperactive lung ILC2s in offspring.

### Prenatal inflammation transcriptionally enhances lung ILC2 hyperactivation programs in neonatal offspring

We next investigated the effects of prenatal inflammation on transcriptional programming of ILC2 hyperactivation *in vivo* by analyzing over 26,000 lung ILC2s (Lin-CD45+IL7rα+Thy1+ST2+) isolated from P14 offspring after poly(I:C) or saline using single-cell RNA sequencing (scRNA-seq). Unsupervised clustering analysis revealed 8 distinct clusters (C1-C8) (**Fig. 3A**), highlighting extensive heterogeneity within our sorted ILC2 population. All clusters expressed canonical ILC2 genes, including *Gata3* (*45–47*)*, Id2* (*34, 46*), *Il7r* and *l1rl1* (*17*) (**Fig. 3B**). Identified clusters were mostly represented across both saline and poly(I:C) conditions (**Fig. 3C**), but to investigate how prenatal inflammation drove ILC2 heterogeneity, we profiled cellular distribution across clusters (**Fig. 3D**). While cell frequency was unchanged among most clusters (C2, C3, C4, C6, C7) in response to prenatal inflammation, C1, C5, and C8 were directly modulated by poly(I:C) (**Fig. 3D**). C1 decreased in cellular density while C5 and C8 were both significantly expanded in response to prenatal inflammation (**Fig. 3D**). C1 was characterized by a gene expression profile found in both ILC2s *(CCR9, CD24a)* (*48, 49*) and ILC3s (*Il22, Klrb1b*) (*50, 51*) (**Fig. 3E)**, a transcriptional signature in line with observed plasticity among ILC populations (*52*). C5 was defined by an activated ILC2 gene signature (*Gata3, Klrg1, Il5, Il13, Il1rl1*) (*20, 53*) (**Fig. 3E)**, while C8, primarily identifiable and expanded under the poly(I:C) condition only, was defined by both a progenitor- (*Il18r1, Zbtb16*) (*54, 55*) and memory-like (*Syne1, Il1r12*) (*53*) gene expression profile (**Fig. 3E**).

**Fig. 3:**
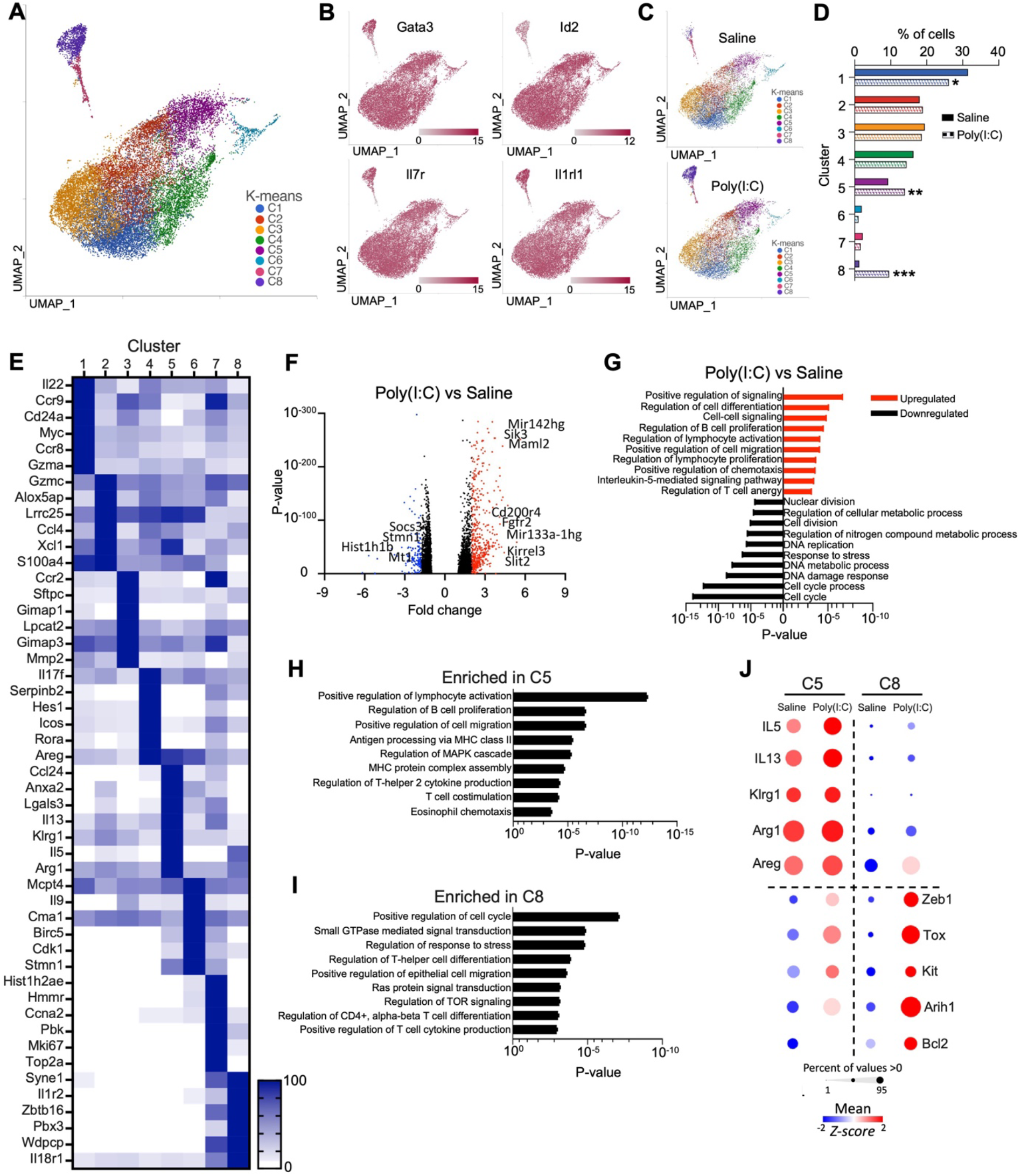
Prenatal inflammation transcriptionally enhances lung ILC2 hyperactivation programs in neonatal offspring. **A)** UMAP of sorted P14 lung ILC2s (CD45+Lin-IL7ra+Thy1+ST2+) across saline and poly(I:C) treated conditions. **B)** UMAPs depicting normalized expression of *Gata3, Id2, Il7r,* and *Il1rl1* (ST2) across saline and poly(I:C) treated conditions. **C)** UMAP representation across saline and poly(I:C) treated conditions. **D)** ILC2 cellular distribution across clusters in response to saline or poly(I:C). **E)** Heat map of differentially expressed genes per cluster across saline or poly(I:C) treated conditions. Legend represents normalized gene expression. **F)** Volcano plot of differentially expressed genes in response to prenatal inflammation. **G)** GSEA pathways based on differentially expressed genes between saline and poly(I:C). P-values are plotted as up- or down-regulated based on poly(I:C) to saline comparison. **H)** GSEA pathways based on differentially expressed genes between saline and poly(I:C) were determined within C5. **I)** GSEA pathways based on differentially expressed genes between saline and poly(I:C) were determined within C8. **J)** Bubble plot comparing expression of ILC2 biomarkers across saline and poly(I:C) for clusters 5 and 8. Bubble size is proportional to the percentage of cells in a cluster expressing that gene and color intensity is proportional to the average scaled gene expression within a cluster (z-score). Data represents average ± SEM. *p ≤ 0.05, **p ≤ 0.01, ***p ≤ 0.001, ****p ≤ 0.0001;

Prenatal inflammation induced global differential overexpression of 507 genes and downregulation of 55 genes compared to saline controls (**Fig. 3F)**. Prenatal inflammation significantly upregulated expression of genes associated with ILC2 hyperactivation, including *Gata3, Il1rl1, Klrg1, Il5, Il13,* and *Bcl2* (**SFig. 1A**). Top upregulated genes included genes associated with asthma risk (*56–58*), including *Maml2, Fgfr2, Stab1, Asxl1, Slmap*, and microRNAs *Mir142hg and Mir133a-hg,* while *Socs3, Stmn1,* and *Mt1* were among the most downregulated genes in response to prenatal inflammation (**Fig. 3F, SFig. 1B**) Pathway analysis on differentially expressed genes across clusters showed increased enrichment for immune activation, specifically IL-5 signaling, as well as proliferation and immune cell recruitment, while being negatively enriched for cell cycling and proliferation (**Fig. 3G**), mirroring our in vivo data (**Fig. 1E-H**). Many of the enriched pathways defining C5 (**Fig. 3H**) mirrored pathways enriched for differentially expressed genes across all clusters (**Fig. 3G**), including immune activation and chemotaxis. C5 was further enriched for MAPK signaling, a canonical signaling pathway downstream of IL-33R, suggesting active Th2-cytokine production (*59, 60*), as well as MHC class-II presentation, indicating their capacity to activate CD4+ Th-cells directly (*61*). Enriched pathways defining C8 included positive regulation of the cell cycle, regulation of stress response and growth signaling pathways, and T-cell differentiation and function. As prenatal inflammation specifically expanded both C5 and C8, we examined differential gene expression in response to poly(I:C) in these two clusters (**Fig. 3J**). Prenatal inflammation increased the expression of activation genes that defined C5, including *Il5, Il13, Klrg1,* and *Arg1* (**Fig. 3J**). In contrast, prenatal inflammation robustly enhanced a stem-like phenotype in C8, as evidenced by increased expression of stem-like genes *Zeb1, Tox, Kit, Arih1,* and *Bcl2* (**Fig. 3J**). Many of these same genes were also upregulated in C5 but to a lesser extent. Together, these data indicate that prenatal inflammation transcriptionally reprograms postnatal ILC2s by expanding specific subsets associated with hyperactivated, inflammatory, or stem-like states.

### ILC2 hyperactivation is accompanied by remodeling of the lung immune landscape

During infection or barrier-disruption, ILC2-derived cytokines play a pivotal role in modulating the early immune response. IL-5 induces recruitment and early activation of eosinophils into the lungs (*25*) while IL-13 is a potent driver of mucus overproduction and smooth muscle contraction (*24, 26*). As we observed that prenatal inflammation induced a hyperactive state in neonatal lung ILC2s both functionally and transcriptionally, we next determined how hyperactive ILC2s impacted the developing lung immune landscape in offspring. Under homeostatic conditions, immune cells present in the P14 lung included B- and T-lymphocytes and a range of myeloid cells, including monocytes, alveolar macrophages, and eosinophils (**Fig. 4A**). Immune cellularity in the lung increased across early postnatal development (**Fig. 4B-H)**, although alveolar macrophages and eosinophils showing a slight decrease at P6 in response to prenatal inflammation. By P14, B-cells, T-cells, NK cells, and eosinophils were significantly expanded in offspring exposed to prenatal inflammation (**Fig. 4A-H).** Notably, in response to prenatal inflammation, ILC2s were expanded as early as P9, before expansion of all other profiled immune cell subsets (**Fig. 4B-H**, **Fig. 1B**). Moreover, ILC2s also displayed the most significant expansion in response to prenatal inflammation, compared to all other immune cells (**Fig. 4A**, **Fig. 1B**). ILC2s both directly and indirectly regulate many of the immune cells that were expanded after prenatal inflammation (*62–64*). Importantly, ILC2s can specifically promote CD4+ Th2 cell activation, which is critical for adaptive immune responses that further enhance B-cell class switching, eosinophil recruitment and the generation of alternately activated macrophages (*27, 60, 65*). We therefore investigated whether early ILC2 expansion and hyperactivation in response to prenatal inflammation correlated with altered T-cell fates within the expanded CD4+ T-cell compartment (**Fig. 4A**). Frequencies of Tbet+ Th1 and Gata3+ Th2 cells within the CD3+CD4+ T-cell fraction were not significantly different under saline conditions. In contrast, we observed sharp skewing of CD3+CD4+ T cells towards a Th2 fate in response to prenatal inflammation, as evidenced by lower frequencies of Th1 cells and a robust increase in the frequency of Th2 cells within the expanded T-cell subset (**Fig. 4I**). Concomitant with Th2 skewing, FOXP3+ T-regulatory cells were also significantly decreased (**Fig. 4J**). These data indicate that prenatal inflammation results in selective remodeling of the lung immune landscape and skewing towards Th2-like immunity correlated with ILC2 expansion and hyperactivation.

**Fig. 4:**
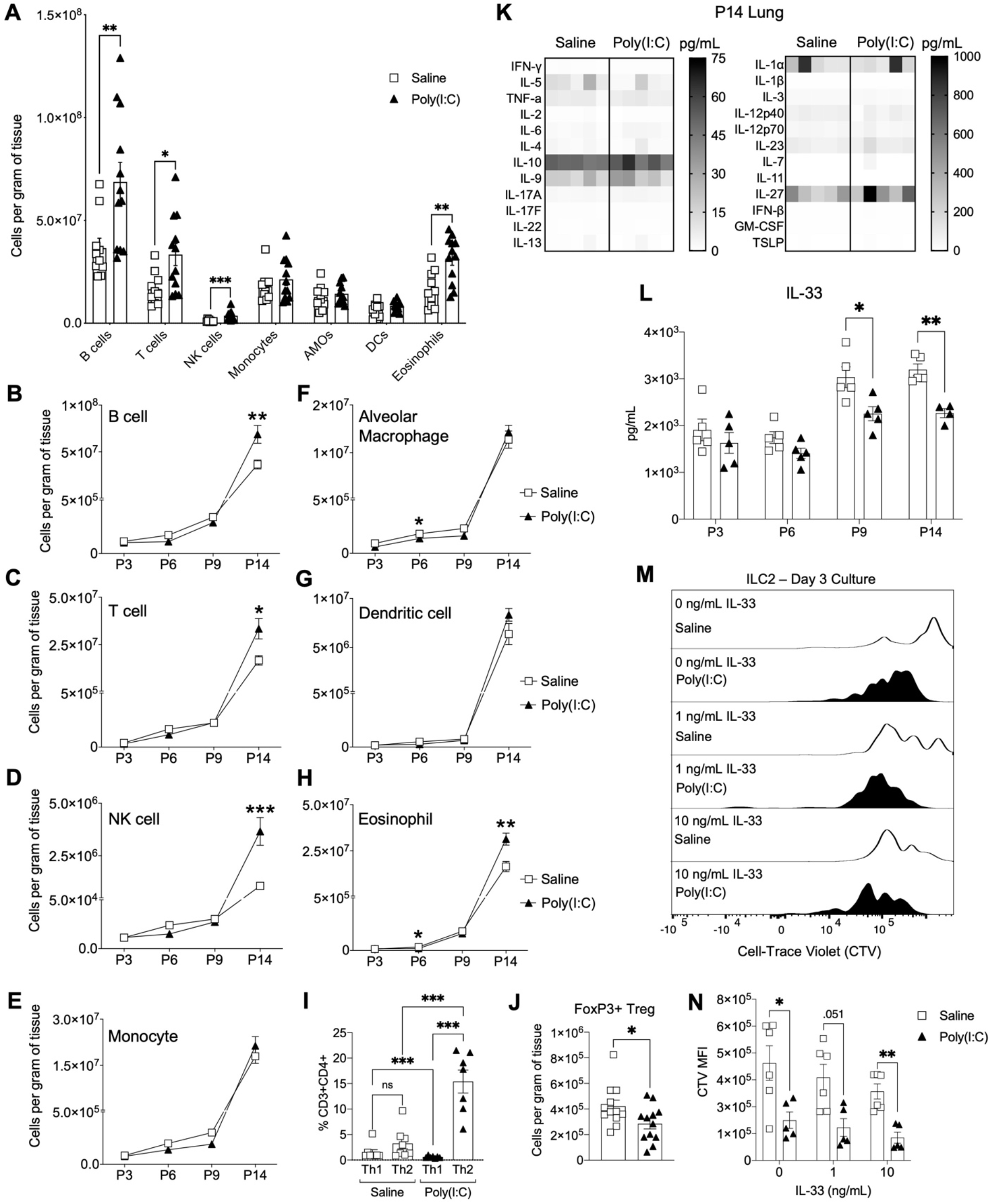
ILC2 expansion is accompanied by remodeling of the lung immune landscape in response to prenatal inflammation. **A)** Quantification of immune populations in the neonatal lung at P14 after saline or poly(I:C). **B-H)** Time course quantification of lung B-cells **(B)** T-cells **(C)** Natural Killer (NK) cells **(D)** Monocytes **(E)** Alveolar Macrophages **(F)** Dendritic cells **(G)** and Eosinophils **(H)** across P3-P14. **I)** Frequency of lung Th1 (Tbet+) or Th2 (Gata3+) cells within the CD3+CD4+ T-cell fraction at P14 after saline or poly(I:C). **J)** Quantification of lung T-regulatory cells (FOXP3+CD25+) at P14 after saline or poly(I:C). **K)** Quantification of cytokines in P14 lung homogenate after saline or poly(I:C). **L)** Quantification of IL-33 cytokine in lung homogenate across P3-P14 from saline or poly(I:C). **M)** Representative histogram depicting cell trace violet (CTV) expression in lung ILC2s sorted from P14 mice after saline or poly(I:C) cultured in 0 ng/mL, 1 ng/mL, or 10 ng/mL IL-33 for 3 days. **N)** MFI of CTV expression in ILC2s after 3-day culture of sorted P14 ILC2s from saline or poly(I:C) treated offspring in 0 ng/mL, 1 ng/mL, or 10 ng/mL IL-33. For all experiments, data represents average ± SEM representing at least two independent experiments. *p ≤ 0.05, **p ≤ 0.01, ***p ≤ 0.001, ****p ≤ 0.0001.

We next examined the effect of prenatal inflammation on the lung cytokine milieu in offspring by quantifying expression levels of 25 different cytokines at P14 (**Fig. 4K-L**). Surprisingly, prenatal inflammation did not alter levels of any profiled cytokines except for IL-33, which was robustly decreased at P14 (**Fig. 4L**). This finding prompted us to profile IL-33 levels across the early postnatal period. IL-33 levels were comparably decreased as early as P9, when ILC2s began to expand in the lung (**Fig. 4L**; **Fig. 1B**). The overall decrease in IL-33 levels in the lung in response to prenatal inflammation conflicted with our observation of expanded and hyperactivated ILC2s, given that epithelial cell-derived IL-33 is known to potently activate ILC2s and induce their rapid expansion and cytokine production. We hypothesized that prenatal inflammation induced a compensatory response in ILC2s associated with hypersensitivity to lower levels of IL-33, as suggested by increased IL-33R (*Il1rl1*) gene expression in ILC2s in response to prenatal inflammation (**SFig. 1A**). To investigate this possibility, we sorted P14 lung ILC2s from saline or prenatal inflammation-exposed offspring and cultured them for 3 days in recombinant IL-33 to measure changes in proliferation capacity using Celltrace Violet^TM^ (CTV) (**Fig. 4M-N**). Across all culture conditions, ILC2s from the prenatal inflammation condition exhibited increased proliferation in response to IL-33 stimulation as compared to saline (**Fig. 4M-N**), indicating a hypersensitive response. Altogether, these data demonstrate that prenatal inflammation induces immune remodeling in a cell-intrinsic manner, whereby ILC2s exhibit adaptive compensatory responses to decreased IL-33 levels in the lung.

### Prenatal inflammation enhances immune response to acute respiratory challenge in neonates

ILC2s have been implicated in the development of allergic airway inflammation (*15, 16, 18, 27*) and elevated numbers of lung ILC2s are associated with more severe outcomes in human allergic asthma patients (*29, 30, 66, 67*). To investigate whether prenatal inflammation-induced expansion of hyperactivated ILC2s increased susceptibility to allergic airway inflammation, we implemented an acute model of allergic inflammation using the allergen papain and measured the response across multiple immune subsets (**Fig. 5A-C, SFig. 2A-E**). At P14, neonatal offspring from litters exposed to saline or prenatal inflammation received 25μg papain or saline intranasally for 4 consecutive days before sacrifice 24 hours later (**Fig. 5A**). Acute papain exposure resulted in significant expansion of ILC2s, B-cells, and dendritic cells in both offspring exposed to prenatal inflammation and saline (**Fig. 5B, SFig. 2A, 2E**), whereas NK cells and alveolar macrophages were unaffected (**SFig. 2C-D**). T-cells responded to papain only in saline-exposed offspring (**SFig. 2B**). ILC2s and eosinophils were the only subsets expanded by exposure to prenatal inflammation alone, prior to papain challenge, and ILC2s also showed the most significant response to papain challenge. Importantly, eosinophils were responsive to acute papain exposure only in offspring exposed to prenatal inflammation (**Fig. 5C**), whereas eosinophils were unresponsive to papain in saline-exposed offspring. Histological analysis of neonatal airways revealed mucus granule accumulation, mucus cell metaplasia, and increased immune cell infiltration along the airways after papain challenge in both saline and prenatal inflammation conditions (**Fig. 5D-H**). These data suggest that ILC2s, activated by prenatal inflammation, drive eosinophil expansion in the lung of offspring that correlate with papain-induced airway inflammation.

**Fig. 5:**
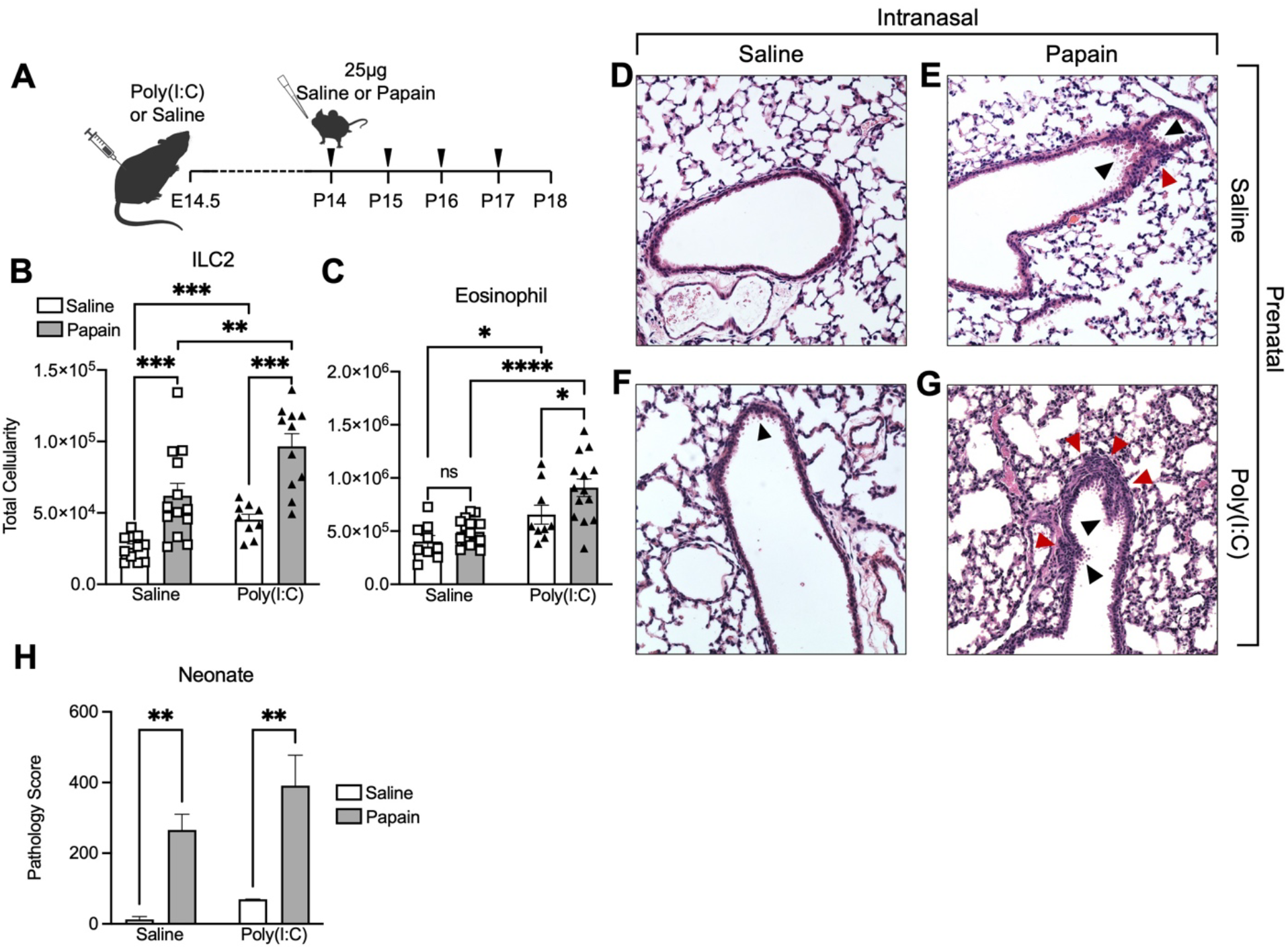
Prenatal inflammation enhances immune response to acute respiratory challenge in neonates. **A)** Schematic of acute neonatal intranasal (i.n.) challenge with papain (25ug) or saline following prenatal saline or poly(I:C) **B-C)** Quantification of ILC2s **(B)** and Eosinophils **(C)** at P18 following i.n. papain (25ug) or saline (**Fig. 5A**) in prenatal saline or poly(I:C) conditions **D-G)** Representative histopathological images of neonatal airways after prenatal inflammation or saline and acute allergen challenge (**Fig. 5A**). Black arrows indicate mucus granule deposition and mucus cell metaplasia. Red arrows indicate immune cell infiltration along the airways. **H)** Pathological scoring in neonatal lungs after i.n. saline or papain challenge in offspring from prenatal saline or poly(I:C). Data represents average ± SEM representing at least two independent experiments. *p ≤ 0.05, **p ≤ 0.01, ***p ≤ 0.001, ****p ≤ 0.0001.

### Prenatal inflammation causes lung dysfunction in adulthood driven by persistent ILC2 expansion in response to acute allergen challenge

We next determined if the effects of prenatal inflammation on lung immune responses to allergen challenge persisted into adulthood. Offspring from litters exposed to prenatal inflammation or saline were aged to adulthood (P70, 10 weeks) before undergoing acute papain challenge (**Fig. 6A**). Overall, as compared to neonatal mice (**Fig. 5**), acute papain challenge did not induce robust changes across most immune cells examined (**SFig. 2F-J**). Prior to allergen challenge, expanded lung ILC2s persisted through adulthood in prenatal inflammation-exposed offspring (**Fig. 6B**), consistent with persistent eosinophilia (**Fig. 6C**). Although ILC2s did not expand in response to papain challenge in saline-exposed adult offspring, there was a trend towards a greater degree of increase in ILC2s in prenatal inflammation-exposed adult offspring in response to papain. The difference in adulthood likely reflects differences in cell number and activation state of ILC2s during adulthood as compared to the neonatal period (*20, 53*). Eosinophils were the only cell type that consistently responded to papain in both saline- and prenatal inflammation-exposed adult offspring (**Fig. 6C**). Histological analysis of airway inflammation confirmed immune infiltration along the airways within the adult lung, as well as mucus cell hyperplasia and mucus granule production in response to papain challenge in both saline and prenatal inflammation conditions (**Fig. 6D-G**). Notably, and in contrast to observations in neonates, adult mice exposed to prenatal inflammation demonstrated significant lung pathology in the absence of papain challenge that was comparable to papain-induced inflammation in saline-exposed offspring (**Fig. 6H**).

**Fig. 6:**
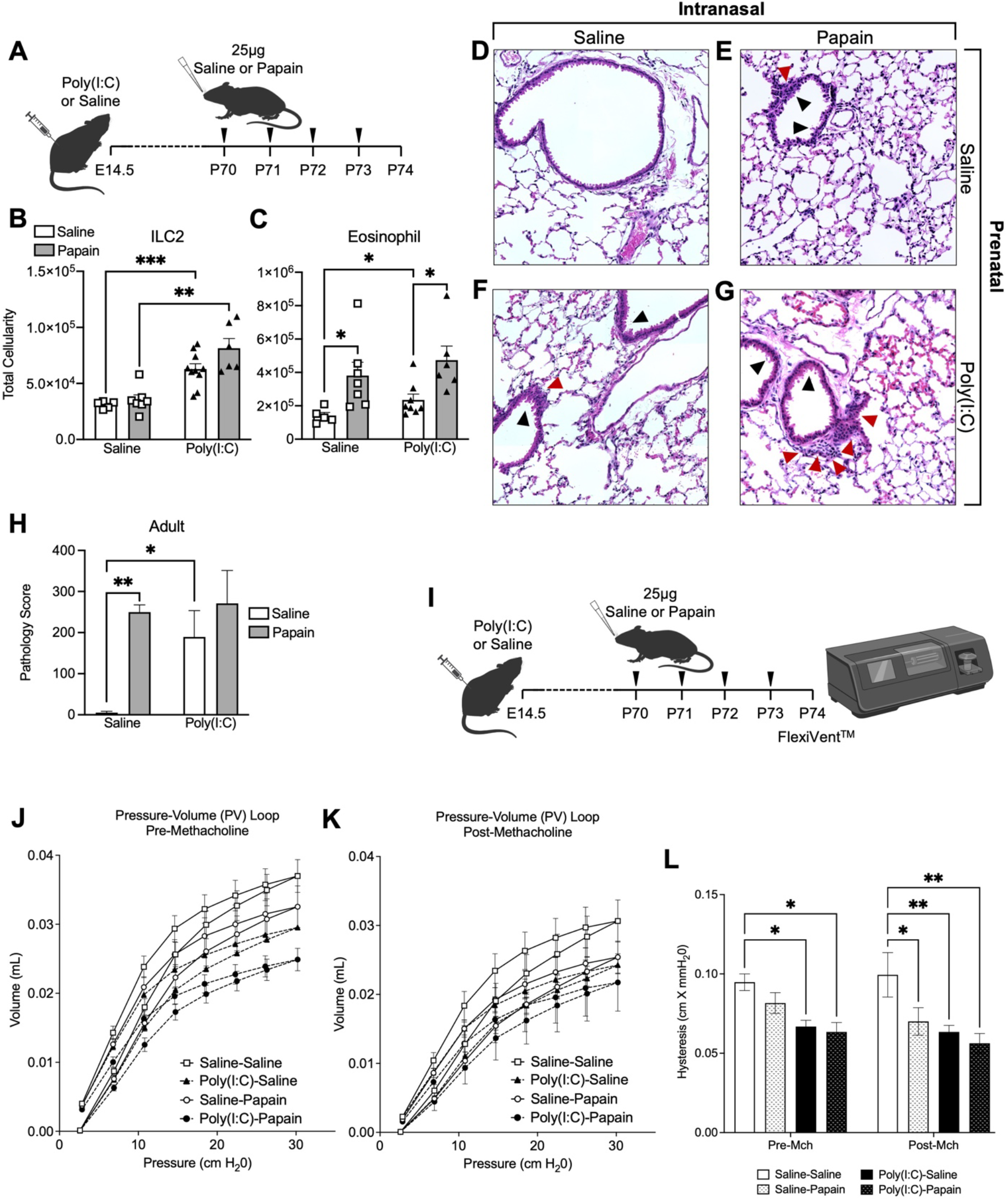
Prenatal inflammation causes lung dysfunction in adulthood driven by persistent ILC2 expansion in response to acute allergen challeng. **A)** Schematic of acute adult intranasal (i.n.) challenge with papain (25ug) or saline following prenatal saline or poly(I:C) **B-C)** Quantification of ILC2s **(B)** and eosinophils **(C)** at P74 following i.n. papain (25ug, **Fig. 6A**) or saline in prenatal saline or poly(I:C) treated conditions **D-G)** Representative histopathological images of adult airways after prenatal inflammation or saline and acute allergen challenge (**Fig. 6A**). Black arrows indicate mucus granule deposition and mucus cell metaplasia. Red arrows indicate immune cell infiltration along the airways. **H)** Pathological scoring in adult lungs after i.n. saline or papain (**Fig. 6A**) in offspring from prenatal saline or poly(I:C) conditions. **I)** Schematic of acute adult i.n. challenge with papain (25ug) or saline, following prenatal saline or poly(I:C), prior to pulmonary mechanical testing using the flexiVent^TM^ system **J-K)** Pre- and post-methacholine Pressure-Volume (PV) loop in adult mice after saline or papain (as shown in **6I**) in offspring from prenatal saline or poly(I:C) conditions. **L)** Quantification of hysteresis (PV-loop area) pre- and post-methacholine challenge of adult mice after saline or papain in offspring from prenatal saline or poly(I:C) conditions. Data represents average ± SEM representing at least two independent experiments. *p ≤ 0.05, **p ≤ 0.01, ***p ≤ 0.001, ****p ≤ 0.0001.

As adult offspring exposed to prenatal inflammation exhibited persistent ILC2 expansion, immune infiltration, and lung pathology (**Fig. 6B-H**), we next assessed changes in pulmonary mechanics using the flexiVent^TM^ system, the gold standard for *in vivo* respiratory mechanic measurements (**Fig. 6I**). To gain insight into the lung’s elastic properties in adult mice, we measured pressure-volume (PV) loops generated from mechanical inflation and deflation of the lungs at baseline and in response to methacholine challenge (**Fig. 6J-K**) and quantified differences as measured by hysteresis (**Fig. 6L**) (*68, 69*). Prior to methacholine challenge, prenatal inflammation alone significantly impaired respiratory function regardless of saline or papain challenge, whereas papain alone had no significant effect on respiratory function in saline-exposed offspring, consistent with worsened histopathology in response to prenatal inflammation alone (**Fig. 6D-H**). Saline-exposed offspring only exhibited respiratory impairment after both methacholine and papain challenge, but prenatal inflammation-exposed offspring demonstrated more severe impairment after methacholine challenge alone. Together, these data indicate that prenatal inflammation causes lung dysfunction through a persistent hyperactivated and expanded ILC2 lung compartment. Our data further indicate that persistent ILC2s act as drivers of eosinophilia and sensitive mediators of allergen-induced airway inflammation.

## DISCUSSION

ILC2s have been implicated as major players in asthma pathogenesis but their role as drivers of asthma susceptibility in early life is unknown. Here we demonstrated that prenatal inflammation programs fetal liver progenitors to produce hyperactive ILC2s that persist into the postnatal lung. Hyperactive ILC2s capable of increased IL-5 and IL-13 production were expanded in the postnatal lung and associated with a remodeled lung immune landscape skewed toward Th2 immunity. Furthermore, prenatal inflammation promoted eosinophilia associated with hyperactive ILC2s and increased susceptibility to allergen-induced airway inflammation and airway dysfunction in adulthood. Our data elucidate a mechanism by which inflammation promotes asthma susceptibility from early-life by skewing the function of tissue-resident ILC2s that seed the developing lung and contribute to the remodeling of tissue immunity.

ILC2s play a pivotal role in tissue homeostasis (*64, 70*), and their potent IL-5 and IL-13 production in response to IL-33 directs both immune cell recruitment and activation during a critical period of postnatal lung development (*21, 22*). In our model, prenatal inflammation induced robust ILC2 hyperactivation, including enhanced proliferation leading to a 3-fold expansion in the postnatal lung, a 2-3-fold increase in stimulated IL-5 and IL-13 production, and increased responsiveness to IL-33. Cumulatively, this multi-faceted functional hyperactivation poised ILC2s to remodel lung immunity as they seeded the developing lung. Consequentially, ILC2 hyperactivation was linked to remodeling of the lung immune landscape in offspring, including Th2 skewing, reduced Tregs, and eosinophilia. Reduced lung IL-33 levels, despite robust activation of ILC2s, provocatively suggests the possibility of rewired immune-epithelial crosstalk as a consequence of ILC2 hyper-responsiveness. Indeed, others have demonstrated that ILC2s can serve as a “sink” for IL-7 in a way that modulates lymphocyte development in the bone marrow (*71*). ILC2 overexpression of IL33R in response to prenatal inflammation could similarly serve as a “sink” for IL-33, thereby regulating immune expansion and activation in the developing lung, including T-regulatory cells. These data emphasize that tissue-resident immune cells that seed developing tissues can play a critical role in instructing tissue immunity. Our data demonstrate that prenatal inflammation or other extrinsic cues can exacerbate the phenotype or function of the earliest infiltrating tissue-resident cells, with major implications for tissue function.

The ILC2-IL-33 axis is a critical response program to lung barrier disruption induced by bacterial, viral and allergen burden (*18, 72*). Papain-induced lung inflammation results in increased IL-33 levels and eosinophilia in ILC2-sufficient *Rag1^-/-^* mice that lack B- and T-cells (*17*), implicating ILC2s as immediate and sufficient initiators of allergen-induced lung inflammation. Similarly, we demonstrated that hyperactivated ILC2s were the main responders to acute intranasal papain challenge and specifically drove eosinophilia in the lung in neonates and adults. Importantly, prenatal inflammation did not cause overt lung pathology in neonates, indicating that prenatal inflammation did not directly affect lung development. Instead, prenatal inflammation alone manifested persistently elevated and hyperactivated ILC2s into adulthood causing sustained eosinophilia, a canonical asthma feature in mice and humans. These persistent cellular changes directly translated into the same airway dysfunction, immune infiltration, and mucus cell metaplasia in adulthood in response to prenatal inflammation alone as that invoked by acute papain challenge. These data highlight the critical effects of early-life inflammation on lung immune health as a driver for sustained lung dysfunction.

Recent work shows that ILC2s display unique phenotypic and transcriptional profiles across ontogeny, suggesting a link between inherent functional heterogeneity and layered immune establishment. Our single-cell analysis directly paralleled our in vivo analysis by revealing that prenatal inflammation expanded and enhanced a hyperactivated ILC2 phenotype, C5, identified by increased *Il5, Il13* and *Klrg1* expression. Transcriptional profiling of neonatal ILC2s also revealed a more hyperactive profile, including increased expression of *Il5*, *Il13*, and *Areg*, as compared to adult ILC2s (*20, 49*). We also identified an expanded stem-like cluster unique to the prenatal inflammation condition, C8, transcriptionally resembling recently identified ILC progenitors in the lung and BM (*73*). This cluster exhibited increased proliferation and enhanced metabolism, suggesting that prenatal inflammation expands a local progenitor in the postnatal lung. In response to prenatal inflammation, lymphocyte activation, chemotaxis, and IL-5 signaling were amongst the most upregulated pathways across all clusters, corroborating in vivo evidence that hyperactivated ILC2s can drive remodeling of the lung immune landscape. These data demonstrate that prenatal inflammation increases lung ILC2 heterogeneity associated with increased asthma risk by 1) generating an ILC2 progenitor-like subset that may serve as a precursor for hyperactivated ILC2s and 2) perturbing layered ILC2 development and expanding pools of hyperactivated “neonatal”-type ILC2s in the lung. Using multiple overlapping fate-mapping models, Schneider et al demonstrated the temporally regulated contribution of fetal, neonatal, and adult ILC2 subsets to the lung, without identifying specific precursors to those subsets (*20*). We specifically demonstrated that the developmentally-restricted HSC (drHSC), a transient, lymphoid-biased fetal progenitor responsible for producing innate-like lymphocytes during fetal development (*41*), is a fetal liver precursor for ILC2s. We further demonstrated that the drHSC produced hyperactivated ILC2s in response to prenatal inflammation upon adoptive transfer. Our findings are therefore consistent with prenatal inflammation influencing layered ontogeny in the postnatal lung by selectively expanding the contribution of one of more early sources of hyperactivated ILC2s, as evidenced by the expansion of both drHSCs (*39*) and CHILPs in the fetal liver by prenatal inflammation. Hyperactivation of the drHSC by prenatal inflammation similarly resulted in the programming of hyperactivated innate-like B1-B cell progeny postnatally and into adulthood (*39*). Both the drHSC and, ultimately, ILC2 expansion are likely driven by a maternal type I interferon response during fetal development, as we also demonstrated that maternal but not fetal deletion of TLR3 ablated drHSC (*39*) and ILC2 expansion.

Our work identifies a fetal ILC2 progenitor that can be programmed during development to produce hyperactive ILC2 progeny in response to prenatal inflammation. This work further contributes to understanding of how perturbation of layered ontogeny of tissue-resident cells in response to prenatal inflammation can contribute to impaired tissue immunity, tissue function, and susceptibility to disease in the postnatal period.

## EXPERIMENTAL MODEL AND SUBJECT DETAILS

### Study design

The goal of this study was to determine how prenatal inflammation altered the establishment and function of ILC2s in offspring to promote allergic airway inflammation. ILC2 development and immune function in offspring was assessed across various timepoints, in the context of prenatal inflammation using viral-mimetic poly(I:C), via *in-vivo* immune profiling, transplantation, *in-vitro* culture, scRNA-seq, histopathology, genetic mouse models of TLR3-deficiency and allergic airway inflammation to assess the role of hyperactive ILC2s on allergic asthma susceptibility.

### Mice

All mice were on a C57BL/6 background and were bred and maintained in-house under specific pathogen-free conditions at the animal facility of the University of California Merced and the University of Utah. Breeders were kept on breeder diet (3980X-081423M) and all other experimental animals were kept on standard diet (2920X-080123M). 8- to 12-week-old female mice were used for experiments and all experiments involving offspring used littermates or independently verified littermate controls. Flkswitch mice (*41, 43, 74*) were generated by crossing Flk2-Cre mice (*75*) to mTmG mice (*76*). For all experiments using Flkswitch mice, WT C57BL/6 females (RRID: IMSR_JAX:000664) were crossed to Flkswitch males as the Flk-Cre transgene is only transmitted through the Y chromosome. TLR3 KO/+ intercrossed mice were generated by crossing TLR3 KO/KO males or females (RRID:IMSR_JAX:005217) to WT C57BL/6 females or males. All experiments were conducted according to Institutional Animal Care and Use Committee (IACUC)-approved protocols.

### Cell isolation

For prenatal inflammation induction, pregnant dams were weighed and injected intraperitoneally with 20mg/kg polyinosinic:polycytidylic acid (poly(I:C), Sigma) or saline Lung cells were isolated from the whole lung using the Miltenyi lung dissociation kit (130-095-927) and gentleMACS^TM^ tissue dissociator with heating blocks. Bone marrow cells were isolated by crushing femurs and tibias in staining media (SM, 1X PBS + 0.5mM EDTA + 2% FBS) using a mortar and pestle and homogenizing marrow plugs using a micropipette. Fetal liver cells were isolated by dissecting whole liver from embryos and mechanically homogenizing the liver in staining media using a micropipette to single-cell suspension. Red blood cell lysis was performed on all samples using 1 mL of 1X ACK (Ammonium-Chloride-Potassium) lysing buffer for one minute, on ice. Reaction was quenched with 10 mLs SM. Samples were filtered through a 70µM filter before antibody staining.

### Antibodies

Biotin-conjugated anti-CD3ε, CD4, CD5, CD8, CD19, NK1.1, Gr1, CD11b, CD11c, F4/80, Ter119, PE-Cy7-conjugated anti-IL7rα, CD64, CD25, CD8, BV480-conjugated anti-Ly6G, BV605-conjugated anti-CD90.2, MHC-II, Tbet, BV785-conjugated anti-KLRG1, CD19, Pacific Blue (PB)-conjugated anti-Sca1, CD4, BV421-conjugated anti-NK1.1, Sca1, PE-conjugated anti-CD11b, IL7rα, IL-13, APC-conjugated anti-ST2, LPAM-1, IL-5, APC-Cy7-conjugated anti-SiglecF, cKit, Alexa Fluor 647-conjugated anti-FOXP3, FITC-conjugated anti-CD3, PE-Cy5-conjugated anti-Flk2 (Flt3), Gata3, were purchased from Biolegend. BUV395-conjugated streptavidin and BUV805-conjugated anti-CD45 were purchased from BD Bioscience. PE-conjugated anti-ST2 was purchased from MD Bioproducts. Propidium iodide or DAPI (BD Bioscience) was used to exclude non-viable cells.

### Cell identification by flow cytometry

All samples were incubated with antibody cocktails containing FC-block (CD16/32-PGP) for 20 mins on ice and protected from light. Lung immune populations were identified as follows: Lineage dump or “Lin” for lung immune populations (CD16/32-PGP, Fcεr1α, CD3, CD4, CD5, CD8, CD19, NK1.1, Gr1, CD11b, CD11c, F4/80, Ter119); ILC2 (Lin-, CD45+, KLRG1+, Sca1+, CD90.2+, ST2+), B cells (CD45+, Ter119-, Ly6G-, CD3-, CD19+), T cells (CD45+, Ter119-, Ly6G-, CD3+, CD19-), NK cells (CD45+, Ter119-, Ly6G-, CD3-, CD19-, CD11c-, MHCII-, CD11b+, CD64-, NK1.1+), Monocytes (CD45+, Ter119-, Ly6G-, CD3-, CD19-, NK1.1-, CD11c-, CD11b+, CD64+), Alveolar Macrophages (CD45+, Ter119-, Ly6G-, CD3-, CD19-, NK1.1-, CD11b^Low/+^+, SiglecF+, CD11c+), Dendritic cells (CD45+, Ter119-, Ly6G-, CD3-, CD19-, NK1.1-, MHCII+, CD11c+), Eosinophils (CD45+, Ter119-, Ly6G-, CD3-, CD19-, NK1.1-, CD11b^Hi^+, SiglecF+, CD11c^Low/-^). For fetal liver and bone marrow populations: CHILP (Lin-, CD45+, IL7rα+, CD25-, Flk2-, α4β7+). BD LSR II and Cytek Aurora were used for phenotypic analysis; BD FACS Aria III and Cytek Aurora CS were used for cell sorting. Flowjo v.10 was used for data analysis.

### Intracellular staining

For dead cell exclusion, samples were incubated in 1X PBS with Zombie Violet™ (Bioegend) or Ghost Dye™ Red 710 (Cytek) fixable viability dye for 20 mins, protected from light, at room temperature. Fixable viability dye was washed off using staining media containing 2% FBS prior to staining with antibody cocktail against cell surface receptors. Quantification of intracellular staining for IL-5 and IL-13 production in lung ILC2s was performed using the Cytofix/Cytoperm kit (BD Biosciences) on 4 x 10^6^ lung cells after 3-hour restimulation with PMA (70ng/mL) and ionomycin (700ng/mL) (Sigma) in 1mL of RPMI-1640 media + 10% heat-inactivated FBS + Brefeldin A (BFA, Biolegend) at 37C. For transcription factor staining, 4 x 10^6^ lung cells were fixed/permeabilized using the True-Nuclear Transcription Factor buffer set (Biolegend) followed by intracellular staining.

### Proliferation of lung ILC2s

Lung cells were dissociated, processed into a single cell suspension, and run through a 70µM filter. Samples were staining with antibody cocktail for ILC2 surface markers and a fixable viability dye. Cells were then fixed and permeabilized with the True-Nuclear Transcription Factor buffer set (Biolegend) and stained intracellularly stained with Ki67-APC (Invitrogen, Carlsbad, CA, USA) and Hoescht 33342 (Invitrogen).

### Fetal HSC transplants

Transplantations were performed by sorting fetal liver GFP+ drHSCs at E15.5 from Flkswitch embryos after maternal poly(I:C) or saline treatment at E14.5. Recipient C57BL/6 mice (8-12 weeks) were sub-lethally irradiated using 600 cGy (Precision X-Rad 320). 200 sorted GFP+ drHSCs wells were transplanted in 1X PBS via retroorbital injection using a 1 mL tuberculin syringe in a volume of 100-200 µL. Lung ILC2 chimerism was determined in recipients at 4 weeks post-transplantation by flow cytometry using the Cytek Aurora.

### ILC2 sort for single-cell RNA sequencing (scRNA-seq)

Pregnant C57BL/6 dams were exposed to poly(I:C) or saline at E14.5 and litters were sacrificed at postnatal day 14. Whole lung is enzymatically digested and mechanically homogenized using Miltenyi lung dissociation kit (130-095-927) in conjunction with a Miltenyi gentleMACS™ tissue dissociator. Red blood cell lysis was performed using 1 mL of 1X ACK (Ammonium-Chloride-Potassium) lysing buffer for one minute, on ice. Reaction was quenched with 10mLs of 1X PBS + 5mM EDTA + 2% FBS (Staining media, SM). Cells were stained with anti-IL7ra biotin antibody on ice, protected from light, for 20 mins. Cells were incubated with Streptavidin MicroBeads (Miltenyi) and run through an LS column (Miltenyi) on magnetic separator (Miltenyi) to enrich for IL7ra+ cells. Enriched cells were stained with ILC2 antibody cocktail. Just before sorting, cells were resuspended in propidium iodide to exclude dead cells via flow cytometry. ILC2s were sorted as Lin-CD45+IL7rα+Thy1+ST2+ at 2,500 events/sec into collection media (SM + 10% FBS). Cells were pelleted after sorting and media swapped to 1X PBS + 0.04% BSA. A sample library was generated using Chromium 10x V3 and run using NovaSeq.

### scRNA-seq data clustering and gene expression

Single-cell libraries from mouse cells were mapped with Cell Ranger’s count pipeline using mm10 genome reference. Using the demultiplexed and aligned count data from the 10X Genomics software CellRanger, we first removed ambient RNA counts from the gene matrix using scAR (*77*), a deep learning-based program from scVI-tools (*78*). We removed low-quality cells using cutoffs for total genes, total transcripts, and mitochondrial percentage per cell. Cells with fewer than 200 genes and more than 3500 genes were discarded. Furthermore, cells with more than 12% unique molecular identifiers (UMIs) from mitochondrial genes were filtered out. We remove doublets using Doublet Detection (DOI: 10.5281/zenodo.2678042). Genes expressed in fewer than five cells were also excluded from further analysis. This filtering step to eliminate low-quality cells and doublets resulted in 13,916 cells in Saline and 14,820 cells in poly(I:C) for downstream analyses. The gene expression matrix was then normalized and scaled for downstream analysis. Principal components analysis (PCA) was performed on the scaled data, and based on the elbow plot, 20 principal components (PCs) were selected for clustering. This dataset was split into two groups, saline and poly(I:C), then analyzed individually using the same steps. K-means clustering revealed eight distinct clusters that were then compared. For transcript normalization, we used scran normalization method (*79*). Even after applying batch effect correction, we did not find significant differences in the cluster annotations of the cells compared with their clustering pattern without batch effect correction. We used the Leiden algorithm (*80*) in conjunction with various forms of dimensionality reduction to identify distinct cell clusters based on RNA expression. We used MAST (*81*) to identify differentially expressed genes for each cluster and treatment. We used GSEA (*82*) for pathway enrichment analyses of transcriptomic data. Finally, we used a scalable denoising application (*83*) to map distinct genes per cluster.

### Quantitation of cytokines in postnatal lung

Neonatal lungs from saline or poly(I:C) treated litters were homogenized in tissue protein extraction reagent (T-PER, Thermo Fisher Scientific) and Complete Mini Protease Inhibitor Cocktail tablets (Sigma) before pelleting to collect supernatant. Supernatant was frozen at −80°C and samples were analyzed on BD Canto (BD Biosciences) following manufacturer recommendations using the Mouse T Helper Cytokine Panel Version 3 and Mouse Cytokine Panel 2 LEGENDplex^TM^ (Biolegend) bead-based immunoassay for the following cytokines: IL-1α, IL-1β, IL-3, IL-12p40, IL-12p70, IL-23, IL-7, IL-11, IL-27, IL-33, IFN-β, GM-CSF, TSLP, IFN-γ, IL-5, TNF-α, IL-2, IL-6, IL-4, IL-10, IL-9, IL-17A, IL-17F, IL-22, and IL-13. Data were analyzed with LEGENDplex^TM^online analysis platform. https://legendplex.qognit.com/.

### ILC2 Culture

ILC2s (Lin-, CD45+, Thy1+, Sca1+) were sorted from lungs of offspring after prenatal saline or poly(I:C) treatment at E14.5. Prior to plating, ILC2s were stained with CellTrace^TM^ Violet (Thermo Fisher Scientific). ILC2s were plated in triplicates at 5,000 cells/well in RPMI-1640 (Sigma) containing 10% FBS, 10 mM HEPES (Sigma), 100 mM non-essential amino acids (Sigma), 1 mM sodium pyruvate (GIBCO), 100 U/mL, 50 mM 2-mercaptoethanol (GIBCO), 10 ng/ml rIL-7 (Peprotec), 10 ng/ml rIL-2 (Peprotec), and 0-10ng/ml rIL-33 (Peprotec) in 96-well round bottom plates at 37C under 5% CO_2_ for 3 days. After 3 days, cells were harvested, stained with ILC2 antibody cocktail, and assessed for CellTrace^TM^ Violet staining via flow cytometry.

### Instransal allergen challenge and pulmonary mechanics assessment

For intranasal allergen challenge, mice were anesthetized by isofluorane inhalation, followed by intranasal injection of Papain allergen (Sigma, 25µg) in 10µls of 1X PBS on days 0, 1, 2, and 3. Mice were sacrificed 24 hours after final saline or papain injection and processed for flow cytometry, histopathology, or pulmonary mechanics assessment. For studies involving respiratory mechanics, mice were anesthetized with ketamine and xylazine (100 + 16 mg/kg, i.p.). When unresponsive, the trachea was surgically cannulated using a 1.2 mm outer diameter, 8mm length tracheal cannula (Harvard apparatus, Holliston, MA) and sutured in place. Mice were then connected to a FlexiVent (SCIREQ, Montreal, PQ, Canada) at positive end-expiratory pressure of 3 cm H2O, and the paralytic vecuronium (2 mg/kg, i.p.) administered. Immediately following, the deep inflation maneuver (inflating the lung from 3 to 40 cmH2O over 3s and then holding at 30 cmH2O for 3s) is performed 2x to ensure proper inflation of the lung prior to pulmonary mechanic measurements. Baseline mouse mechanics were performed before and after serial delivery of methacholine using the FlexiVent Aeroneb fine particle nebulizer (0, 3.125, 6.25, 12.5, 25, and 50 mg/mL) in saline for 10s per dose at 4-5 min intervals. The quasistatic pressure volume curve (PV loop) was generated in 7 equal steps between 3 and 30 cm H2O and from which hysteresis (area of PV loop) was calculated. Mice undergoing lung function studies were not used for histological, immunohistochemical, or flow cytometric analysis. Data were normalized to mouse weight.

### Histopathology

Mice were perfused using 10% of neutral buffered formalin and intubated intratracheally to inflate lungs. Lungs were fixed for 48 hours at room temperature. Samples were paraffin embedded and sections were stained for hematoxylin and eosin (H&E). Mouse lungs were cleared of blood by cardiac perfusion and whole lungs were fixed by tracheal instillation using 10% neutral buffered formalin. Following paraffin embedding, sections were cut and stained with hematoxylin and eosin (H&E) by the Associated Regional and University Pathologists (ARUP) Inc., at the University of Utah. Histological scoring was performed by a blinded pathologist as follows: (% of bronchial/bronchiolar epithelium with infiltrate x measured number of cellular depth of peribronchial infiltrate) + (% of pulmonary veins with infiltrate x measured number of cellular depth of perivenous infiltrate). This score was calculated on 2 slides per animal and 2–4 animals per group. Representative images were taken on a Zeiss Axiovert 200 microscope with an Optronics Microfire camera. Images are shown in Fig. 5 and Fig. 6 at 20X magnification.

### Statistical Analysis

Prism v8.0 (GraphPad Software, San Diego, CA, USA) was used to perform statistical tests. Mice used were randomized across age, sex, and cage for any given experiment. Statistically significant differences between groups with similar variance as determined by the SEM and SD were assessed by either un-paired two-tailed t-tests for parametric data, or Mann–Whitney tests for non-parametric data. Šidák’s multiple comparisons test was used for comparisons between prenatal inflammation or saline treated litters after saline or papain injection pre- and post-methacholine challenge (FlexiVent). P-values were calculated and are indicated as follows: *p ≤ 0.05; **p ≤ 0.01; ***p ≤ 0.001; ****p ≤ 0.0001. Sample sizes are indicated in each figure and unless otherwise noted, all experiments were performed in at least two independent replicates.

## Author Contributions

Conceptualization: AEB, DAL

Methodology: AEB, DAL, RSW, CD-R, KJW

Investigation: DAL, CD-R, AG, LM-A

Visualization: DAL, RSW, CD-R, EJM, KJW, AEB

Funding acquisition: AEB

Project administration: AEB

Supervision: AEB, DAL

Writing – original draft: AEB, DAL

Writing – review & editing: AEB, DAL

## Data and materials availability

All data are available in the main text or the supplementary materials. Data for scseq have been deposited in XXX with https://www.ncbi.nlm.nih.gov/geo/info/submission.html.

## Supplementary Materials

### Supplementary Figures

**Supplementary Figure 1:**
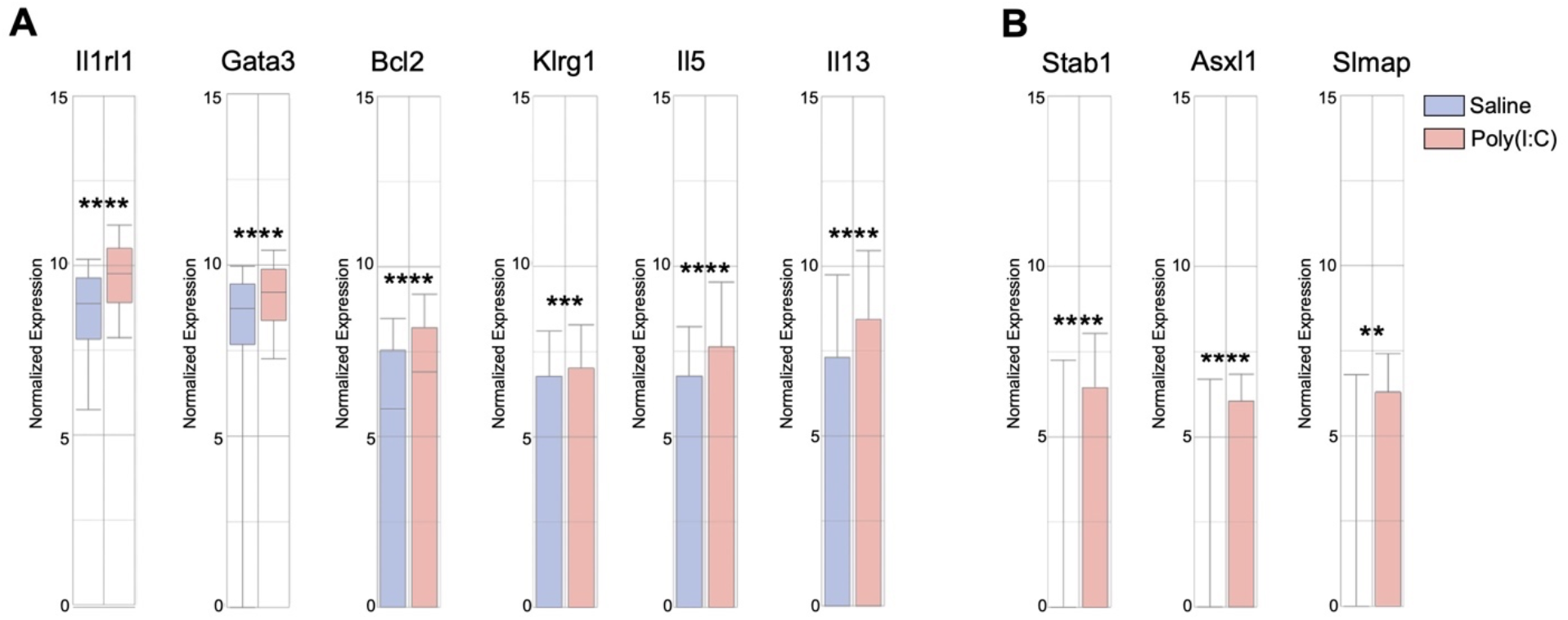
Prenatal inflammation upregulates genes associated with ILC2 identity and genome-wide association studies (GWAS) of asthma. **A)** Violin plots showing normalized expression of genes associated with ILC2 identity across all clusters in saline or poly(I:C) treated conditions. **B)** Violin plots showing normalized expression of genes identified in GWAS of asthma risk across all clusters in saline or poly(I:C) treated conditions. Data represents average ± SD representing at least two independent experiments.. *p ≤ 0.05, **p ≤ 0.01, ***p ≤ 0.001, ****p ≤ 0.0001; Omission of statistical demarcation indicates differences are not statistically significant.

**Supplementary Figure 2:**
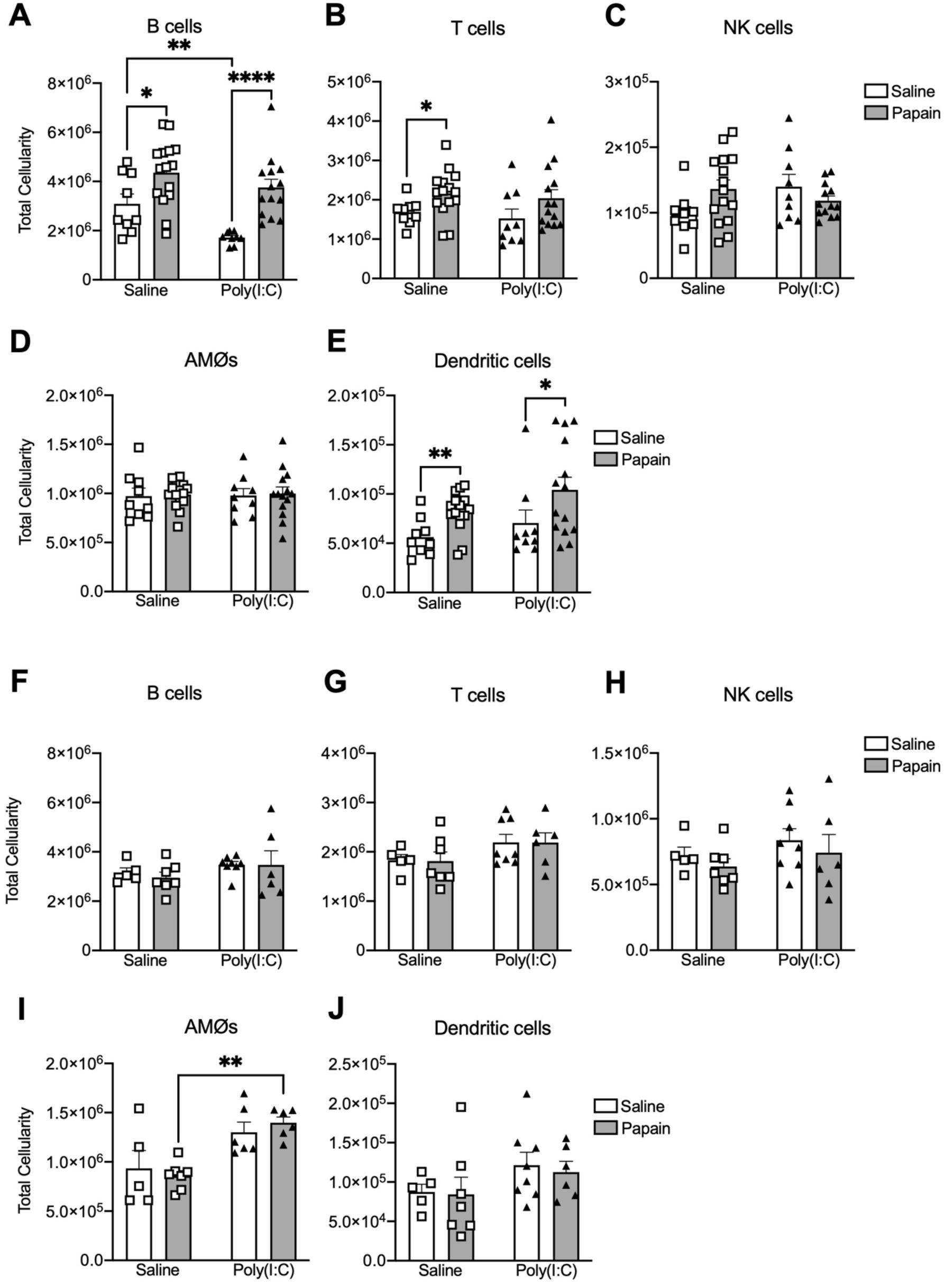
Prenatal inflammation induces a differential response of lung immune cells to papain challenge in neonatal and adult offsprin. **A-E)** Quantification of B-cells **(A),** T-cells **(B),** NK cells **(C),** AMØs **(D),** and Dendritic cells **(E)** at P18 following i.n. injection of papain (25ug, as shown in **Fig. 5A**) or saline in prenatal saline or poly(I:C) treated conditions. **F-J)** Quantification of B-cells **(F),** T-cells **(G),** NK cells **(H),** AMØs **(I),** and Dendritic cells **(J)** at 10 weeks following i.n. injection of papain (25ug, as shown in **Fig. 6A**) or saline in prenatal saline or poly(I:C) treated conditions. Data represents average ± SEM representing at least two independent experiments. *p ≤ 0.05, **p ≤ 0.01, ***p ≤ 0.001, ****p ≤ 0.0001;

## References

1. M. Korppi, A. Kotaniemi-Syrjanen, Infection-induced wheezing during the first year of life does not mean asthma: a 50-year-old observation. Acta Paediatr 92, 1494–1495 (2003).

2. N. Sigurs, P. M. Gustafsson, R. Bjarnason, F. Lundberg, S. Schmidt, F. Sigurbergsson, B. Kjellman, Severe respiratory syncytial virus bronchiolitis in infancy and asthma and allergy at age 13. Am J Respir Crit Care Med 171, 137–141 (2005).

3. R. T. Stein, D. Sherrill, W. J. Morgan, C. J. Holberg, M. Halonen, L. M. Taussig, A. L. Wright, F. D. Martinez, Respiratory syncytial virus in early life and risk of wheeze and allergy by age 13 years. Lancet 354, 541–545 (1999).

4. P. G. Holt, P. D. Sly, Viral infections and atopy in asthma pathogenesis: new rationales for asthma prevention and treatment. Nat Med 18, 726–735 (2012).

5. P. Wu, W. D. Dupont, M. R. Griffin, K. N. Carroll, E. F. Mitchel, T. Gebretsadik, T. V. Hartert, Evidence of a causal role of winter virus infection during infancy in early childhood asthma. Am J Respir Crit Care Med 178, 1123–1129 (2008).

6. M. K. Hyvarinen, A. Kotaniemi-Syrjanen, T. M. Reijonen, K. Korhonen, M. O. Korppi, Teenage asthma after severe early childhood wheezing: an 11-year prospective follow-up. Pediatr Pulmonol 40, 316–323 (2005).

7. D. J. Jackson, M. D. Evans, R. E. Gangnon, C. J. Tisler, T. E. Pappas, W. M. Lee, J. E. Gern, R. F. Lemanske, Jr., Evidence for a causal relationship between allergic sensitization and rhinovirus wheezing in early life. Am J Respir Crit Care Med 185, 281–285 (2012).

8. J. K. Yoo, T. S. Kim, M. M. Hufford, T. J. Braciale, Viral infection of the lung: host response and sequelae. J Allergy Clin Immunol 132, 1263–1276; quiz 1277 (2013).

9. S. Trouillet-Assant, S. Viel, A. Ouziel, L. Boisselier, P. Rebaud, R. Basmaci, N. Droz, A. Belot, S. Pons, K. Brengel-Pesce, Y. Gillet, E. Javouhey, G. Antoine Study, Type I Interferon in Children with Viral or Bacterial Infections. Clin Chem 66, 802–808 (2020).

10. J. E. Mold, J. M. McCune, Immunological tolerance during fetal development: from mouse to man. Advances in immunology 115, 73–111 (2012).

11. A. C. Apostol, K. D. C. Jensen, A. E. Beaudin, Training the Fetal Immune System Through Maternal Inflammation-A Layered Hygiene Hypothesis. Front Immunol 11, 123 (2020).

12. J. V. Fahy, Type 2 inflammation in asthma--present in most, absent in many. Nat Rev Immunol 15, 57–65 (2015).

13. S. Basha, N. Surendran, M. Pichichero, Immune responses in neonates. Expert Rev Clin Immunol 10, 1171–1184 (2014).

14. G. P. Tsafaras, P. Ntontsi, G. Xanthou, Advantages and Limitations of the Neonatal Immune System. Front Pediatr 8, 5 (2020).

15. K. R. Bartemes, K. Iijima, T. Kobayashi, G. M. Kephart, A. N. McKenzie, H. Kita, IL-33-responsive lineage-CD25+ CD44(hi) lymphoid cells mediate innate type 2 immunity and allergic inflammation in the lungs. J Immunol 188, 1503–1513 (2012).

16. M. J. Gold, F. Antignano, T. Y. Halim, J. A. Hirota, M. R. Blanchet, C. Zaph, F. Takei, K. M. McNagny, Group 2 innate lymphoid cells facilitate sensitization to local, but not systemic, TH2-inducing allergen exposures. J Allergy Clin Immunol 133, 1142–1148 (2014).

17. T. Y. Halim, A. MacLaren, M. T. Romanish, M. J. Gold, K. M. McNagny, F. Takei, Retinoic-acid-receptor-related orphan nuclear receptor alpha is required for natural helper cell development and allergic inflammation. Immunity 37, 463–474 (2012).

18. R. G. Klein Wolterink, A. Kleinjan, M. van Nimwegen, I. Bergen, M. de Bruijn, Y. Levani, R. W. Hendriks, Pulmonary innate lymphoid cells are major producers of IL-5 and IL-13 in murine models of allergic asthma. Eur J Immunol 42, 1106–1116 (2012).

19. J. L. Barlow, A. Bellosi, C. S. Hardman, L. F. Drynan, S. H. Wong, J. P. Cruickshank, A. N. McKenzie, Innate IL-13-producing nuocytes arise during allergic lung inflammation and contribute to airways hyperreactivity. J Allergy Clin Immunol 129, 191–198.e191-194 (2012).

20. C. Schneider, J. Lee, S. Koga, R. R. Ricardo-Gonzalez, J. C. Nussbaum, L. K. Smith, S. A. Villeda, H. E. Liang, R. M. Locksley, Tissue-Resident Group 2 Innate Lymphoid Cells Differentiate by Layered Ontogeny and In Situ Perinatal Priming. Immunity 50, 1425–1438.e1425 (2019).

21. I. M. de Kleer, M. Kool, M. J. de Bruijn, M. Willart, J. van Moorleghem, M. J. Schuijs, M. Plantinga, R. Beyaert, E. Hams, P. G. Fallon, H. Hammad, R. W. Hendriks, B. N. Lambrecht, Perinatal Activation of the Interleukin-33 Pathway Promotes Type 2 Immunity in the Developing Lung. Immunity 45, 1285–1298 (2016).

22. S. Saluzzo, A. D. Gorki, B. M. J. Rana, R. Martins, S. Scanlon, P. Starkl, K. Lakovits, A. Hladik, A. Korosec, O. Sharif, J. M. Warszawska, H. Jolin, I. Mesteri, A. N. J. McKenzie, S. Knapp, First-Breath-Induced Type 2 Pathways Shape the Lung Immune Environment. Cell Rep 18, 1893–1905 (2017).

23. M. Duchesne, I. Okoye, P. Lacy, Epithelial cell alarmin cytokines: Frontline mediators of the asthma inflammatory response. Front Immunol 13, 975914 (2022).

24. N. K. Malavia, J. D. Mih, C. B. Raub, B. T. Dinh, S. C. George, IL-13 induces a bronchial epithelial phenotype that is profibrotic. Respir Res 9, 27 (2008).

25. M. M. Fort, J. Cheung, D. Yen, J. Li, S. M. Zurawski, S. Lo, S. Menon, T. Clifford, B. Hunte, R. Lesley, T. Muchamuel, S. D. Hurst, G. Zurawski, M. W. Leach, D. M. Gorman, D. M. Rennick, IL-25 induces IL-4, IL-5, and IL-13 and Th2-associated pathologies in vivo. Immunity 15, 985–995 (2001).

26. D. A. Kuperman, X. Huang, L. L. Koth, G. H. Chang, G. M. Dolganov, Z. Zhu, J. A. Elias, D. Sheppard, D. J. Erle, Direct effects of interleukin-13 on epithelial cells cause airway hyperreactivity and mucus overproduction in asthma. Nat Med 8, 885–889 (2002).

27. T. Y. Halim, C. A. Steer, L. Matha, M. J. Gold, I. Martinez-Gonzalez, K. M. McNagny, A. N. McKenzie, F. Takei, Group 2 innate lymphoid cells are critical for the initiation of adaptive T helper 2 cell-mediated allergic lung inflammation. Immunity 40, 425–435 (2014).

28. B. Roediger, R. Kyle, S. S. Tay, A. J. Mitchell, H. A. Bolton, T. V. Guy, S. Y. Tan, E. Forbes-Blom, P. L. Tong, Y. Koller, E. Shklovskaya, M. Iwashima, K. D. McCoy, G. Le Gros, B. Fazekas de St Groth, W. Weninger, IL-2 is a critical regulator of group 2 innate lymphoid cell function during pulmonary inflammation. J Allergy Clin Immunol 136, 1653–1663 e1657 (2015).

29. S. G. Smith, R. Chen, M. Kjarsgaard, C. Huang, J. P. Oliveria, P. M. O’Byrne, G. M. Gauvreau, L. P. Boulet, C. Lemiere, J. Martin, P. Nair, R. Sehmi, Increased numbers of activated group 2 innate lymphoid cells in the airways of patients with severe asthma and persistent airway eosinophilia. J Allergy Clin Immunol 137, 75–86 e78 (2016).

30. P. Nagakumar, L. Denney, L. Fleming, A. Bush, C. M. Lloyd, S. Saglani, Type 2 innate lymphoid cells in induced sputum from children with severe asthma. J Allergy Clin Immunol 137, 624–626 e626 (2016).

31. I. E. Ishizuka, S. Chea, H. Gudjonson, M. G. Constantinides, A. R. Dinner, A. Bendelac, R. Golub, Single-cell analysis defines the divergence between the innate lymphoid cell lineage and lymphoid tissue-inducer cell lineage. Nat Immunol 17, 269–276 (2016).

32. Q. Yang, F. Li, C. Harly, S. Xing, L. Ye, X. Xia, H. Wang, X. Wang, S. Yu, X. Zhou, M. Cam, H. H. Xue, A. Bhandoola, TCF-1 upregulation identifies early innate lymphoid progenitors in the bone marrow. Nat Immunol 16, 1044–1050 (2015).

33. C. Harly, M. Cam, J. Kaye, A. Bhandoola, Development and differentiation of early innate lymphoid progenitors. J Exp Med 215, 249–262 (2018).

34. W. Xu, D. E. Cherrier, S. Chea, C. Vosshenrich, N. Serafini, M. Petit, P. Liu, R. Golub, J. P. Di Santo, An Id2(RFP)-Reporter Mouse Redefines Innate Lymphoid Cell Precursor Potentials. Immunity 50, 1054–1068 e1053 (2019).

35. C. S. N. Klose, M. Flach, L. Mohle, L. Rogell, T. Hoyler, K. Ebert, C. Fabiunke, D. Pfeifer, V. Sexl, D. Fonseca-Pereira, R. G. Domingues, H. Veiga-Fernandes, S. J. Arnold, M. Busslinger, I. R. Dunay, Y. Tanriver, A. Diefenbach, Differentiation of type 1 ILCs from a common progenitor to all helper-like innate lymphoid cell lineages. Cell 157, 340–356 (2014).

36. M. G. Constantinides, B. D. McDonald, P. A. Verhoef, A. Bendelac, A committed precursor to innate lymphoid cells. Nature 508, 397–401 (2014).

37. G. Gasteiger, X. Fan, S. Dikiy, S. Y. Lee, A. Y. Rudensky, Tissue residency of innate lymphoid cells in lymphoid and nonlymphoid organs. Science 350, 981–985 (2015).

38. H. Aegerter, B. N. Lambrecht, C. V. Jakubzick, Biology of lung macrophages in health and disease. Immunity 55, 1564–1580 (2022).

39. D. A. López, A. C. Apostol, E. J. Lebish, C. H. Valencia, M. C. Romero-Mulero, P. Pavlovich, G. E. Hernandez, E. C. Forsberg, N. Cabezas-Wallscheid, A. E. Beaudin, Prenatal inflammation perturbs murine fetal hematopoietic development and causes persistent changes to postnatal immunity. Cell Rep 41, (2022).

40. M. Salimi, J. L. Barlow, S. P. Saunders, L. Xue, D. Gutowska-Owsiak, X. Wang, L. C. Huang, D. Johnson, S. T. Scanlon, A. N. McKenzie, P. G. Fallon, G. S. Ogg, A role for IL-25 and IL-33-driven type-2 innate lymphoid cells in atopic dermatitis. J Exp Med 210, 2939–2950 (2013).

41. A. E. Beaudin, S. W. Boyer, J. Perez-Cunningham, G. E. Hernandez, S. C. Derderian, C. Jujjavarapu, E. Aaserude, T. MacKenzie, E. C. Forsberg, A Transient Developmental Hematopoietic Stem Cell Gives Rise to Innate-like B and T Cells. Cell stem cell 19, 768–783 (2016).

42. A. E. Beaudin, S. W. Boyer, E. C. Forsberg, Flk2/Flt3 promotes both myeloid and lymphoid development by expanding non-self-renewing multipotent hematopoietic progenitor cells. Exp Hematol 42, 218–229 e214 (2014).

43. S. W. Boyer, A. E. Beaudin, E. C. Forsberg, Mapping stem cell differentiation pathways from hematopoietic stem cells using Flk2/Flt3L lineage tracing. Cell Cycle 11, 3180–3188 (2012).

44. J. K. Bando, H. E. Liang, R. M. Locksley, Identification and distribution of developing innate lymphoid cells in the fetal mouse intestine. Nat Immunol 16, 153–160 (2015).

45. J. Mjosberg, J. Bernink, K. Golebski, J. J. Karrich, C. P. Peters, B. Blom, A. A. te Velde, W. J. Fokkens, C. M. van Drunen, H. Spits, The transcription factor GATA3 is essential for the function of human type 2 innate lymphoid cells. Immunity 37, 649–659 (2012).

46. T. Hoyler, C. S. Klose, A. Souabni, A. Turqueti-Neves, D. Pfeifer, E. L. Rawlins, D. Voehringer, M. Busslinger, A. Diefenbach, The transcription factor GATA-3 controls cell fate and maintenance of type 2 innate lymphoid cells. Immunity 37, 634–648 (2012).

47. H. Spits, D. Artis, M. Colonna, A. Diefenbach, J. P. Di Santo, G. Eberl, S. Koyasu, R. M. Locksley, A. N. McKenzie, R. E. Mebius, F. Powrie, E. Vivier, Innate lymphoid cells--a proposal for uniform nomenclature. Nat Rev Immunol 13, 145–149 (2013).

48. J. Zhang, J. Qiu, W. Zhou, J. Cao, X. Hu, W. Mi, B. Su, B. He, J. Qiu, L. Shen, Neuropilin-1 mediates lung tissue-specific control of ILC2 function in type 2 immunity. Nat Immunol 23, 237–250 (2022).

49. C. A. Steer, L. Matha, H. Shim, F. Takei, Lung group 2 innate lymphoid cells are trained by endogenous IL-33 in the neonatal period. JCI Insight 5, (2020).

50. L. Van Maele, C. Carnoy, D. Cayet, S. Ivanov, R. Porte, E. Deruy, J. A. Chabalgoity, J. C. Renauld, G. Eberl, A. G. Benecke, F. Trottein, C. Faveeuw, J. C. Sirard, Activation of Type 3 innate lymphoid cells and interleukin 22 secretion in the lungs during Streptococcus pneumoniae infection. J Infect Dis 210, 493–503 (2014).

51. M. Pokrovskii, J. A. Hall, D. E. Ochayon, R. Yi, N. S. Chaimowitz, H. Seelamneni, N. Carriero, A. Watters, S. N. Waggoner, D. R. Littman, R. Bonneau, E. R. Miraldi, Characterization of Transcriptional Regulatory Networks that Promote and Restrict Identities and Functions of Intestinal Innate Lymphoid Cells. Immunity 51, 185–197 e186 (2019).

52. A. A. Korchagina, S. A. Shein, E. Koroleva, A. V. Tumanov, Transcriptional control of ILC identity. Front Immunol 14, 1146077 (2023).

53. I. Martinez-Gonzalez, L. Matha, C. A. Steer, M. Ghaedi, G. F. Poon, F. Takei, Allergen-Experienced Group 2 Innate Lymphoid Cells Acquire Memory-like Properties and Enhance Allergic Lung Inflammation. Immunity 45, 198–208 (2016).

54. M. Ghaedi, Z. Y. Shen, M. Orangi, I. Martinez-Gonzalez, L. Wei, X. Lu, A. Das, A. Heravi-Moussavi, M. A. Marra, A. Bhandoola, F. Takei, Single-cell analysis of RORalpha tracer mouse lung reveals ILC progenitors and effector ILC2 subsets. J Exp Med 217, (2020).

55. Y. Yu, J. C. Tsang, C. Wang, S. Clare, J. Wang, X. Chen, C. Brandt, L. Kane, L. S. Campos, L. Lu, G. T. Belz, A. N. McKenzie, S. A. Teichmann, G. Dougan, P. Liu, Single-cell RNA-seq identifies a PD-1(hi) ILC progenitor and defines its development pathway. Nature 539, 102–106 (2016).

56. S. E. Reese, C. J. Xu, H. T. den Dekker, M. K. Lee, S. Sikdar, C. Ruiz-Arenas, S. K. Merid, F. I. Rezwan, C. M. Page, V. Ullemar, P. E. Melton, S. S. Oh, I. V. Yang, K. Burrows, C. Soderhall, D. D. Jima, L. Gao, R. Arathimos, L. K. Kupers, M. Wielscher, P. Rzehak, J. Lahti, C. Laprise, A. M. Madore, J. Ward, B. D. Bennett, T. Wang, D. A. Bell, B. consortium, J. M. Vonk, S. E. Haberg, S. Zhao, R. Karlsson, E. Hollams, D. Hu, A. J. Richards, A. Bergstrom, G. C. Sharp, J. F. Felix, M. Bustamante, O. Gruzieva, R. L. Maguire, F. Gilliland, N. Baiz, E. A. Nohr, E. Corpeleijn, S. Sebert, W. Karmaus, V. Grote, E. Kajantie, M. C. Magnus, A. K. Ortqvist, C. Eng, A. H. Liu, I. Kull, V. W. V. Jaddoe, J. Sunyer, J. Kere, C. Hoyo, I. Annesi-Maesano, S. H. Arshad, B. Koletzko, B. Brunekreef, E. B. Binder, K. Raikkonen, E. Reischl, J. W. Holloway, M. R. Jarvelin, H. Snieder, N. Kazmi, C. V. Breton, S. K. Murphy, G. Pershagen, J. M. Anto, C. L. Relton, D. A. Schwartz, E. G. Burchard, R. C. Huang, W. Nystad, C. Almqvist, A. J. Henderson, E. Melen, L. Duijts, G. H. Koppelman, S. J. London, Epigenome-wide meta-analysis of DNA methylation and childhood asthma. J Allergy Clin Immunol 143, 2062–2074 (2019).

57. M. C. Altman, M. A. Gill, E. Whalen, D. C. Babineau, B. Shao, A. H. Liu, B. Jepson, R. S. Gruchalla, G. T. O’Connor, J. A. Pongracic, C. M. Kercsmar, G. K. Khurana Hershey, E. M. Zoratti, C. C. Johnson, S. J. Teach, M. Kattan, L. B. Bacharier, A. Beigelman, S. M. Sigelman, S. Presnell, J. E. Gern, P. J. Gergen, L. M. Wheatley, A. Togias, W. W. Busse, D. J. Jackson, Transcriptome networks identify mechanisms of viral and nonviral asthma exacerbations in children. Nat Immunol 20, 637–651 (2019).

58. K. Tsuo, W. Zhou, Y. Wang, M. Kanai, S. Namba, R. Gupta, L. Majara, L. L. Nkambule, T. Morisaki, Y. Okada, B. M. Neale, I. Global Biobank Meta-analysis, M. J. Daly, A. R. Martin, Multi-ancestry meta-analysis of asthma identifies novel associations and highlights the value of increased power and diversity. Cell Genom 2, 100212 (2022).

59. J. Schmitz, A. Owyang, E. Oldham, Y. Song, E. Murphy, T. K. McClanahan, G. Zurawski, M. Moshrefi, J. Qin, X. Li, D. M. Gorman, J. F. Bazan, R. A. Kastelein, IL-33, an interleukin-1-like cytokine that signals via the IL-1 receptor-related protein ST2 and induces T helper type 2-associated cytokines. Immunity 23, 479–490 (2005).

60. A. S. Mirchandani, R. J. Salmond, F. Y. Liew, Interleukin-33 and the function of innate lymphoid cells. Trends Immunol 33, 389–396 (2012).

61. C. J. Oliphant, Y. Y. Hwang, J. A. Walker, M. Salimi, S. H. Wong, J. M. Brewer, A. Englezakis, J. L. Barlow, E. Hams, S. T. Scanlon, G. S. Ogg, P. G. Fallon, A. N. McKenzie, MHCII-mediated dialog between group 2 innate lymphoid cells and CD4(+) T cells potentiates type 2 immunity and promotes parasitic helminth expulsion. Immunity 41, 283–295 (2014).

62. L. Maggi, G. Montaini, A. Mazzoni, B. Rossettini, M. Capone, M. C. Rossi, V. Santarlasci, F. Liotta, O. Rossi, O. Gallo, R. De Palma, E. Maggi, L. Cosmi, S. Romagnani, F. Annunziato, Human circulating group 2 innate lymphoid cells can express CD154 and promote IgE production. J Allergy Clin Immunol 139, 964–976 e964 (2017).

63. L. Y. Drake, K. Iijima, K. Bartemes, H. Kita, Group 2 Innate Lymphoid Cells Promote an Early Antibody Response to a Respiratory Antigen in Mice. J Immunol 197, 1335–1342 (2016).

64. J. C. Nussbaum, S. J. Van Dyken, J. von Moltke, L. E. Cheng, A. Mohapatra, A. B. Molofsky, E. E. Thornton, M. F. Krummel, A. Chawla, H. E. Liang, R. M. Locksley, Type 2 innate lymphoid cells control eosinophil homeostasis. Nature 502, 245–248 (2013).

65. G. Pelaia, A. Vatrella, M. T. Busceti, L. Gallelli, C. Calabrese, R. Terracciano, R. Maselli, Cellular mechanisms underlying eosinophilic and neutrophilic airway inflammation in asthma. Mediators Inflamm 2015, 879783 (2015).

66. T. Liu, J. Wu, J. Zhao, J. Wang, Y. Zhang, L. Liu, L. Cao, Y. Liu, L. Dong, Type 2 innate lymphoid cells: A novel biomarker of eosinophilic airway inflammation in patients with mild to moderate asthma. Respir Med 109, 1391–1396 (2015).

67. R. Chen, S. G. Smith, B. Salter, A. El-Gammal, J. P. Oliveria, C. Obminski, R. Watson, P. M. O’Byrne, G. M. Gauvreau, R. Sehmi, Allergen-induced Increases in Sputum Levels of Group 2 Innate Lymphoid Cells in Subjects with Asthma. Am J Respir Crit Care Med 196, 700–712 (2017).

68. K. J. Warren, C. Deering-Rice, T. Huecksteadt, S. Trivedi, A. Venosa, C. Reilly, K. Sanders, F. Clayton, T. A. Wyatt, J. A. Poole, N. M. Heller, D. Leung, R. Paine, 3rd, Steady-state estradiol triggers a unique innate immune response to allergen resulting in increased airway resistance. Biol Sex Differ 14, 2 (2023).

69. M. Boucher, C. Henry, F. Khadangi, A. Dufour-Mailhot, S. Tremblay-Pitre, L. Fereydoonzad, D. Brunet, A. Robichaud, Y. Bosse, Effects of airway smooth muscle contraction and inflammation on lung tissue compliance. Am J Physiol Lung Cell Mol Physiol 322, L294–L304 (2022).

70. A. B. Molofsky, F. Van Gool, H. E. Liang, S. J. Van Dyken, J. C. Nussbaum, J. Lee, J. A. Bluestone, R. M. Locksley, Interleukin-33 and Interferon-gamma Counter-Regulate Group 2 Innate Lymphoid Cell Activation during Immune Perturbation. Immunity 43, 161–174 (2015).

71. C. E. Martin, D. S. Spasova, K. Frimpong-Boateng, H. O. Kim, M. Lee, K. S. Kim, C. D. Surh, Interleukin-7 Availability Is Maintained by a Hematopoietic Cytokine Sink Comprising Innate Lymphoid Cells and T Cells. Immunity 47, 171–182 e174 (2017).

72. D. R. Neill, S. H. Wong, A. Bellosi, R. J. Flynn, M. Daly, T. K. Langford, C. Bucks, C. M. Kane, P. G. Fallon, R. Pannell, H. E. Jolin, A. N. McKenzie, Nuocytes represent a new innate effector leukocyte that mediates type-2 immunity. Nature 464, 1367–1370 (2010).

73. P. Zeis, M. Lian, X. Fan, J. S. Herman, D. C. Hernandez, R. Gentek, S. Elias, C. Symowski, K. Knopper, N. Peltokangas, C. Friedrich, R. Doucet-Ladeveze, A. M. Kabat, R. M. Locksley, D. Voehringer, M. Bajenoff, A. Y. Rudensky, C. Romagnani, D. Grun, G. Gasteiger, In Situ Maturation and Tissue Adaptation of Type 2 Innate Lymphoid Cell Progenitors. Immunity 53, 775–792 e779 (2020).

74. S. W. Boyer, A. V. Schroeder, S. Smith-Berdan, E. C. Forsberg, All hematopoietic cells develop from hematopoietic stem cells through Flk2/Flt3-positive progenitor cells. Cell stem cell 9, 64–73 (2011).

75. C. Benz, V. C. Martins, F. Radtke, C. C. Bleul, The stream of precursors that colonizes the thymus proceeds selectively through the early T lineage precursor stage of T cell development. J Exp Med 205, 1187–1199 (2008).

76. M. D. Muzumdar, B. Tasic, K. Miyamichi, L. Li, L. Luo, A global double-fluorescent Cre reporter mouse. Genesis 45, 593–605 (2007).

77. C. Sheng, R. Lopes, G. Li, S. Schuierer, A. Waldt, R. Cuttat, S. Dimitrieva, A. Kauffmann, E. Durand, G. G. Galli, G. Roma, A. d. Weck, Probabilistic machine learning ensures accurate ambient denoising in droplet-based single-cell omics. bioRxiv, 2022.2001.2014.476312 (2022).

78. A. Gayoso, R. Lopez, G. Xing, P. Boyeau, V. Valiollah Pour Amiri, J. Hong, K. Wu, M. Jayasuriya, E. Mehlman, M. Langevin, Y. Liu, J. Samaran, G. Misrachi, A. Nazaret, O. Clivio, C. Xu, T. Ashuach, M. Gabitto, M. Lotfollahi, V. Svensson, E. da Veiga Beltrame, V. Kleshchevnikov, C. Talavera-Lopez, L. Pachter, F. J. Theis, A. Streets, M. I. Jordan, J. Regier, N. Yosef, A Python library for probabilistic analysis of single-cell omics data. Nat Biotechnol 40, 163–166 (2022).

79. A. T. Lun, K. Bach, J. C. Marioni, Pooling across cells to normalize single-cell RNA sequencing data with many zero counts. Genome Biol 17, 75 (2016).

80. V. A. Traag, L. Waltman, N. J. van Eck, From Louvain to Leiden: guaranteeing well-connected communities. Sci Rep 9, 5233 (2019).

81. G. Finak, A. McDavid, M. Yajima, J. Deng, V. Gersuk, A. K. Shalek, C. K. Slichter, H. W. Miller, M. J. McElrath, M. Prlic, P. S. Linsley, R. Gottardo, MAST: a flexible statistical framework for assessing transcriptional changes and characterizing heterogeneity in single-cell RNA sequencing data. Genome Biol 16, 278 (2015).

82. A. Subramanian, P. Tamayo, V. K. Mootha, S. Mukherjee, B. L. Ebert, M. A. Gillette, A. Paulovich, S. L. Pomeroy, T. R. Golub, E. S. Lander, J. P. Mesirov, Gene set enrichment analysis: a knowledge-based approach for interpreting genome-wide expression profiles. Proc Natl Acad Sci U S A 102, 15545–15550 (2005).

83. G. Eraslan, L. M. Simon, M. Mircea, N. S. Mueller, F. J. Theis, Single-cell RNA-seq denoising using a deep count autoencoder. Nat Commun 10, 390 (2019).

